# MYRF drives heterochronic miRNAs and LIN-42, and amplifies oscillatory programs for stage transitions

**DOI:** 10.64898/2026.01.27.701908

**Authors:** Zhao Wang, Shiqian Shen, Xiaoting Feng, Di Chen, Qian Bian, Yingchuan B. Qi

## Abstract

Larval development in *C. elegans* proceeds through discrete stages, yet the mechanisms that drive stage transitions remain incompletely defined. Here, we identify the transmembrane transcription factor MYRF as a master temporal regulator essential for larval progression. MYRF cleavage and nuclear accumulation oscillate across larval stages, peaking during mid-to-late intermolt phases. MYRF directly activates key heterochronic microRNAs—including *lin-4*, *mir-48/241/84*, and *let-7*—and amplifies the oscillatory gene network by promoting central regulators such as *nhr-23*. MYRF loss-of-function or acute depletion suppresses and flattens this network, causing developmental arrest during the late intermolt phase of each larval cycle and stalling molting, seam cell division, and vulval morphogenesis. Notably, MYRF oscillations precede and induce activation of *lin-42/Period*, which feeds back to repress *myrf*, sharpening oscillatory precision. Together, these findings establish MYRF as a central driver of larval stage transitions, coupling microRNA-conferred temporal identity with an amplifier-based logic that executes stage- and phase-specific developmental programs.

## Introduction

Across animal development, progression through discrete stages is orchestrated by temporally gated transcriptional programs^[1,2]^. Some of the most conspicuous examples of stage transitions include insect metamorphosis^[3]^, which transforms larvae into adults through hormone-triggered waves of gene expression that remodel nearly every tissue, and mammalian puberty^[4]^, which marks a major developmental transition involving coordinated changes across the brain, endocrine system, and reproductive organs. Despite their significance, we still lack a clear understanding of the molecular drivers behind stage transitions.

In *Caenorhabditis elegans*, post-embryonic development progresses through four larval stages (L1–L4), each accompanied by a molt that marks the transition to the next stage^[5]^. Classical genetic studies uncovered the heterochronic pathway, a regulatory hierarchy that establishes stage-specific temporal identity, most prominently through microRNAs such as *lin-4*, *let-7*, and the *let-7* family^[6–10]^. The microRNA *lin-4* is activated in late L1 and promotes L2 cell fate by downregulating the L1-stage determinant LIN-14^[6,9]^. Subsequently, *let-7* family members *mir-48*, *mir-241*, and *mir-84* act around the L2-L3 transition by downregulating L2-stage regulators such as *hbl-1*^[8]^. Additionally, *let-7* functions primarily at the L4-to-adult transition to repress targets including *lin-41* and thereby permit adult-specific gene expression^[10,11]^. These microRNAs together act in a sequential network to confer temporal identity and drive orderly developmental progression. A key unresolved question in this paradigm is what molecular mechanisms trigger the activation of these stage-specific microRNAs. We previously reported that the transmembrane transcription factor MYRF is essential for *lin-4* activation in late L1,^[12]^ but how transcription of *let-7* family microRNAs is regulated has remained unknown.

Molting is a defining feature of each larval stage transition in *C. elegans*^[13–17]^. Molting comprises apolysis (separation of the old cuticle), synthesis of a new cuticle, and ecdysis (shedding of the old cuticle). Entry into molting is accompanied by a behavioral quiescent state known as lethargus. While many structural and enzymatic components of the molting machinery have been identified—including cuticle collagens, proteases, and astacin metalloproteases—the regulatory logic that controls the initiation, coordination, and timing of this multi-step process remains poorly understood. Unlike insects, where steroid hormone signaling plays a central role in regulating molting and metamorphosis^[18]^, no equivalent endocrine pathway has been clearly defined in *C. elegans*^[19]^.

An important temporal regulator of larval molting is *lin-42*, the *C. elegans* homolog of the circadian clock gene *Period*^[20–23]^. *lin-42* expression oscillates with a peak in the late phase of each larval stage, coinciding with the molting time window. Loss of *lin-42* leads to desynchronized molting cycles, molting defects, precocious hypodermal differentiation, and, in some cases, larval arrest. Despite its central role, the upstream mechanisms that drive the oscillatory expression of *lin-42* remain unknown.

Genome-wide transcriptomic analyses have further revealed that more than 3,700 genes exhibit rhythmic expression during larval development, forming a large oscillatory gene network that initiates in late embryogenesis and persists until adulthood^[24]^. These oscillations are aligned with the four molting cycles and peak at defined phases of each larval stage. While many oscillatory genes encode cuticle components or molting enzymes, their functional diversity suggests roles extending beyond molting^[25]^. Several transcription factors have been implicated in regulating subsets of this network, including *nhr-23*, *blmp-1*, *grh-1*, *bed-3*, and *myrf-1*^[26]^. Among these, *nhr-23*, the homolog of mammalian ROR nuclear receptors, is a core component of the molting cycle oscillator^[26–29]^. *nhr-23* exhibits rhythmic expression and directly regulates numerous cuticle genes. However, the regulatory architecture that initiates, sustains, and resets this developmental oscillator remains only partially understood.

*myrf-1(pqn-47)* was first identified in a genetic screen for molting-defective mutants conducted by the Ruvkun laboratory^[30]^, and was later independently identified in our laboratory in a screen for defects in DD motor neuron synaptic remodeling^[31]^, a developmental event tightly coupled to the L1–L2 transition. MYRF is a transmembrane transcription factor that undergoes self-cleavage to release an N-terminal fragment (N-MYRF), which translocates to the nucleus to regulate gene expression^[32–34]^. MYRF is conserved across metazoans; in vertebrates, MYRF is a master regulator of oligodendrocyte myelination, and its loss causes embryonic lethality, while haploinsufficiency in humans leads to congenital developmental disorders^[35,36]^. In *C. elegans*, two paralogs, *myrf-1* and *myrf-2*, are ubiquitously expressed^[31]^. Full-length MYRF localizes to the cell membrane during early and mid L1, and cleavage is temporally regulated, peaking in late L1 and coinciding with key developmental events including *lin-4* activation, DD synaptic remodeling, and molting^[12,31,37]^. *myrf-1* null mutants exhibit penetrant L1 molting defects and perish trapped in unshed cuticles, whereas *myrf-2* null mutants show no obvious growth defects^[30,31]^. Nevertheless, *myrf-1* and *myrf-2* together contribute to *lin-4* activation and DD synaptic remodeling, with *myrf-1* playing a predominant role. Despite the stage-defining significance of heterochronic microRNAs, mutations in these genes—including *lin-4*—do not block larval progression or molting. Therefore, the developmental arrest and molting defects observed in *myrf-1* mutants must arise from additional essential functions of MYRF beyond heterochronic microRNA regulation.

*myrf-1* itself exhibits transcriptional oscillation across all four larval stages^[24]^, and RNAi-mediated depletion of *myrf-1* causes molting defects at multiple stages^[26,30]^, suggesting that MYRF functions broadly throughout larval development rather than at a single transition point. However, how MYRF contributes to the oscillatory gene network and how its function is deployed across individual larval stages have not been definitively established.

In this study, we characterize MYRF as a central regulator of *C. elegans* larval stage transitions. By combining genetic mutants with temporally inducible and tissue-specific degradation approaches, we dissect MYRF-1 and MYRF-2 function across all larval stages. We find that MYRF-1 and MYRF-2 together are required for each stage transition; in their absence, animals progress to the late intermolt phase and then undergo developmental arrest in broad, yet selective somatic tissues, while germline growth continues. Despite these wide-ranging developmental consequences, ChIP-seq analysis reveals that MYRF directly targets a remarkably focused set of genes, including *lin-4*, *let-7*, the *let-7* family microRNAs, and key timing regulators such as *lin-42* and *nhr-23*. Our data reveal that MYRF integrates two core timing systems—heterochronic microRNAs for stage identity and the phase-specific oscillatory network—positioning it as a master regulator of larval stage transition.

## Results

### The Oscillatory Pattern of MYRF-1 Expression and Cleavage

Genome-wide analyses have identified *myrf-1* as part of a large cohort of genes exhibiting oscillatory expression across larval development^[24]^. *myrf-1* mRNA levels display robust cyclic patterns, peaking at the late phase of each intermolt period (Figure 1A). To monitor protein dynamics, we used a previously generated knock-in strain in which GFP is inserted into the N-terminal domain of *myrf-1*, thereby labeling both full-length (membrane-tethered) and cleaved N-terminal (nuclear) MYRF-1^[31,37]^. Imaging revealed a cycle of subcellular localization: membrane-localized full-length MYRF-1 predominates during early to mid-intermolt phases, followed by a nuclear enrichment of cleaved MYRF-1 in the late phase (Figure 1A). As animals enter lethargus, nuclear signals fade, and membrane localization re-emerges with the next cycle.

**Figure 1.**
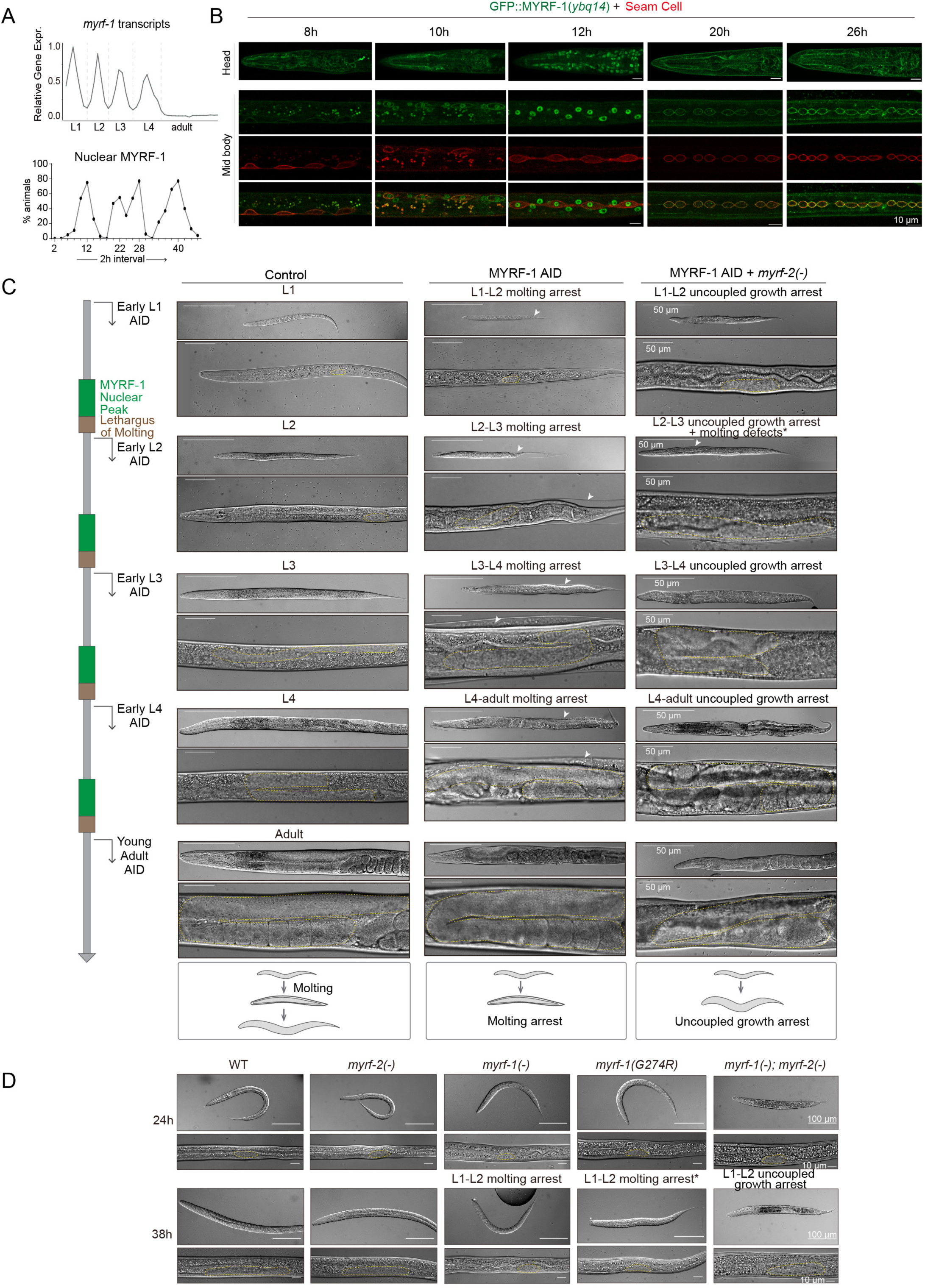
Oscillatory MYRF activity is required for larval stage transitions and molting progression. **A.** Oscillatory expression of *myrf-1* transcripts across larval development (top), based on datasets from Meeuse *et al.*, and corresponding oscillation of nuclear GFP::MYRF-1 levels across a population (bottom), with one peak per larval stage. **B.** Imaging of GFP::MYRF-1 (green) together with seam cell membrane markers (red) reveals cyclic membrane-to-nuclear translocation of MYRF-1 from mid-L1 to mid-L2. Nuclear MYRF-1 peaks coincide with seam cell elongation during mid-to-late intermolt phases and decline during lethargus, whereas membrane-associated MYRF-1 is prominent during the two rounds of seam cell division. **C.** Temporal auxin-induced degradation (AID) of MYRF-1 initiated at defined larval stages; the experimental scheme is shown on the left. Acute MYRF-1 depletion causes stage-specific arrest at the subsequent molt. MYRF-1 AID alone results in molting arrest with unshed cuticle, whereas MYRF-1 AID in a *myrf-2(–)* background leads to uncoupled growth arrest without ecdysis. One exception is MYRF-1 AID; *myrf-2(–)* animals from early L2, which display an uncoupled growth arrest phenotype accompanied by unshed cuticle(*). Gonads are highlighted by yellow dotted outlines. **D.** Developmental phenotypes of wild type and *myrf* mutants at 24 h and 38 h post-hatching. *myrf-1(–)* mutants arrest with L1-L2 molting defects, *myrf-1(G274R)* mutants exhibit extra growth with delayed L1-L2 molting defects(*), and *myrf-1(–); myrf-2(–)* animals exhibit uncoupled growth arrest without molting.

To align these oscillations with developmental events, we used seam cell markers to stage animals (Figure 1B). Nuclear MYRF-1 peaks during seam cell elongation in mid-to-late-intermolt, while membrane-tethered MYRF-1 coincides with seam cell division during molt and early-intermolt, suggesting a precise temporal control linked to cell division and morphogenesis.

### MYRF-1 Is Required for Molting at Every Larval Transition

Because *myrf-1* null mutants arrest at the end of the L1 stage with a characteristic double-cuticle molting defect, the requirement for MYRF beyond the first larval transition has remained unclear. Is MYRF-1 a stage-specific trigger acting only at L1–L2, or a recurrent regulator required at every larval transition? To address this question, we employed an auxin-inducible degradation (AID) system^[38,39]^ to achieve temporally controlled inactivation of MYRF-1 (Figure S1A).

Using CRISPR/Cas9-mediated genome editing, we inserted a Degron::GFP cassette at residue A172 of *myrf-1*, the same position used in the functional *gfp::myrf-1(ybq14)* allele. This *myrf-1::degron::gfp* allele did not cause detectable developmental defects. The degradation was driven by ubiquitously expressed TIR1 in a single-copy transgene. Upon auxin treatment, GFP signals—both membrane-associated and nuclear—disappeared within ∼2 hours in most animals, indicating efficient MYRF-1 degradation (Figure S1A). When gravid adults were placed on auxin-containing plates, their progeny - hatched and grew under AID treatment - uniformly arrested at the L1 molt with unshed cuticle, phenocopying *myrf-1* null mutants and confirming the specificity and effectiveness of the AID system.

To determine whether MYRF-1 is required beyond L1, we synchronized animals to the very early phase of each larval stage by identifying the lethargus of the preceding stage and then induced MYRF-1 degradation. In all cases, animals arrested uniformly at the end of the current stage with molting failure and unshed cuticle (Figure 1C). Dauer larvae treated with MYRF-1 AID exited dauer upon feeding but arrested at the dauer–L4 molt with unshed cuticle (Figure S1B, C). In contrast, animals treated during L4 lethargus progressed to fertile adults without obvious defects (Figure 1C). These results demonstrate that MYRF-1 is required at every larval transition and indispensable for complete molting.

### MYRF-1 Acts during a Late Intermolt Time Window

To define the temporal window of MYRF-1 action, we performed auxin-shift experiments, transferring synchronized larvae to auxin plates at hourly intervals (Figure S1D). MYRF-1 depletion initiated during the early phase of a larval stage caused arrest at the end of that same stage. In contrast, depletion during the late intermolt phase allowed animals to complete the current stage but resulted in arrest at the subsequent molt. Given that nuclear MYRF peaks during late intermolt, these data indicate that MYRF activity during this late window is required to license the upcoming molting transition; once this window has passed, MYRF-1 is dispensable until the next stage.

### MYRF-1/2 Couple Growth to Larval Stage Progression

Strikingly, when MYRF-1 was depleted from early L1 in *myrf-2(-)* background (hereafter referred to as MYRF-deficient), animals exhibited a distinct phenotype from MYRF-1 AID alone (Figure 1C). These animals continued to grow to an apparent L2-like size but then arrested without signs of molting or molting defects. Notably, these arrested animals remained viable on plates for over a week. Although superficially larger, developmental marker analysis revealed that they were in fact arrested at the end of L1 (see section “MYRF Deficiency Stalls the Molting Program”). Because ecdysis was never initiated, no “double cuticle” was observed. These animals continued to grow from late L1 onward, exhibiting pronounced gonadal expansion indicative of ongoing germline proliferation, until eventual arrest. We term this phenotype “uncoupled growth arrest”, characterized by continued size increase and germline growth in the absence of molting or lineage progression. In contrast, the arrest observed in MYRF-1 AID alone—associated with failed ecdysis and double cuticles—is referred to as “molting arrest”.

The late-L1 uncoupled growth arrest phenotype in *myrf*-deficient animals closely resembles that of *myrf-1(-); myrf-2(-)* double-null mutants, whereas the molting arrest observed in MYRF-1 AID alone resembles *myrf-1* single-null mutants (Figure 1D). We previously characterized the *myrf-1(ju1121[G274R])* allele^[31]^, which disrupts MYRF-1 DNA binding and dominantly interferes with MYRF-2 through hetero-oligomerization. *myrf-1(ju1121)* animals grow to an L2-like size but show limited gonad expansion and arrest later than *myrf-1* null mutants, uniformly displaying double cuticles. This *ju1121* phenotype appears intermediate between *myrf-1* single-null and MYRF double-null conditions.

When AID was initiated in early L2, MYRF-deficient animals again displayed uncoupled growth arrest and grew larger than animals experiencing L2 molting arrest from MYRF-1 AID alone (Figure 1C). However, these animals also exhibited defective L2 molting, with old cuticles often detached along the trunk but remaining attached at the head and tail (Figure S1E). This pattern is distinct from the classic “corset” molting defect caused by *nekl-3* loss, in which mid-body cuticle remains constricted^[40]^. Uncoupled growth arrest was similarly observed when MYRF depletion was initiated in early L3 or early L4. In early L3-depleted animals, no molting occurred and vulval invagination—a hallmark of L4 development—was absent (Figure 1C). In early L4-depleted animals, animals grew to adult size and developed oocytes, yet vulval morphogenesis halted at mid-L4, with toroid formation but no eversion (Figure 1C; Figure S1F).

Together, these results demonstrate that MYRF functions as a recurrent and essential regulator of larval stage transitions. MYRF activity during late intermolt is required to execute the upcoming molt and coordinate progression of somatic developmental programs. In the absence of MYRF, animals become locked in a pre-molt state while selective tissues—most notably the germline—continue to grow, revealing an unexpected uncoupling between growth and developmental timing. Thus, MYRF acts as a critical gatekeeper that synchronizes organismal growth with larval stage progression.

### MYRF-Deficiency Stalls the Molting Program

To further define the developmental arrest caused by MYRF inactivation, we analyzed cuticular identity and molting progression using stage-specific markers. We leveraged a recently developed systematic toolkit of endogenously tagged cuticular collagens that provides a robust molecular definition of cuticle identity across larval stages, enabling direct discrimination between molting initiation and cuticle replacement defects^[41]^. Using this framework, we focused on collagen genes specifically expressed in L1 or L2 stages to assess cuticle synthesis and shedding. *dpy-14*, an L1-specific collagen gene, incorporates into the L1 cuticle and is released during L1 ecdysis (Figure 2A). In *myrf-1* AID or single mutants, animals exhibited a “double cuticle” phenotype, with outer DPY-14-positive cuticle retained and inner cuticle lacking DPY-14. This indicates molting initiation occurred, and the new cuticle lacked L1 identity. In contrast, *myrf*-deficient or *myrf-1/2* double mutants retained DPY-14 fluorescence throughout (Figure 2A), and shed cuticles were never observed on the plate, indicating a complete failure to initiate ecdysis. Tracking single animals confirmed that in wild-type animals, shed DPY-14-labeled cuticles are found on the plate, while in *myrf-*deficient animals, no shedding occurred (Figure S2A).

**Figure 2.**
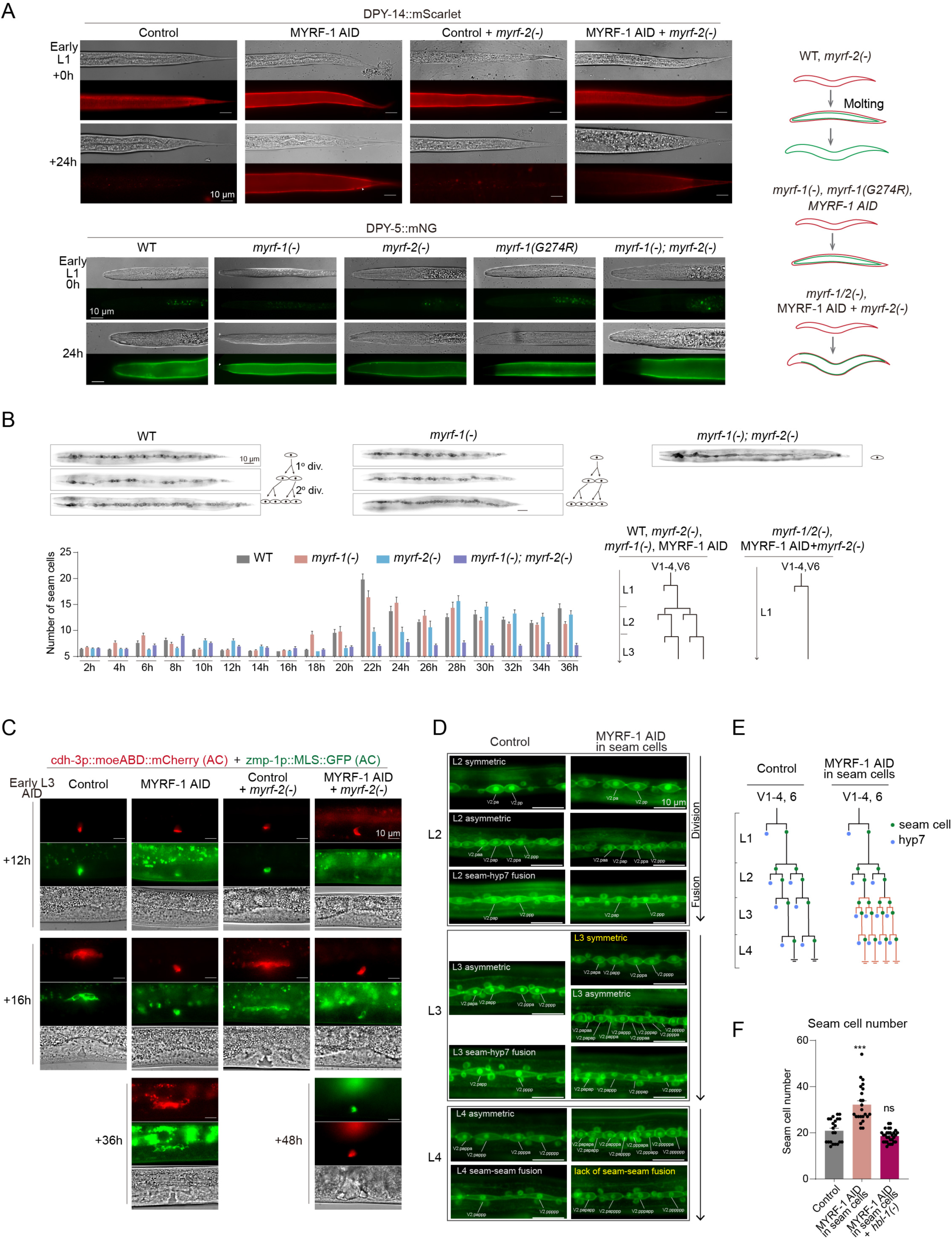
MYRF is required for cuticle remodeling, seam cell patterning, and late larval developmental programs. **A.** Analysis of L1 cuticle shedding using stage-specific collagen reporters. DPY-14::mScarlet (top) labels the L1 cuticle, and DPY-5::mNeonGreen (bottom) labels the L2 cuticle. Control and *myrf-2(–)* animals shed the DPY-14–marked L1 cuticle normally. *myrf-1* loss-of-function animals retain partially shed DPY-14–positive cuticles, whereas *myrf-1/2*–deficient animals retain DPY-14 signal without detectable shedding. Right, schematic summary of molting outcomes. **B.** Seam cell division analysis. Representative images and quantification (bottom) show that WT, *myrf-2(–)*, and *myrf-1(–)* animals undergo two rounds of seam cell divisions during the L1–L2 transition, whereas seam cell divisions are absent in *myrf-1; myrf-2* double mutants. Right, schematic summary of seam cell lineages. **C.** Anchor cell (AC) and vulval development visualized using *cdh-3p* and *zmp-1p* reporters. MYRF-1 AID animals complete AC–UTSE fusion with a delay relative to controls, whereas MYRF-deficient animals fail to execute this process. **D.** Seam cell–specific MYRF-1 depletion reveals reiteration of L2-like symmetric divisions during L3. Subsequent asymmetric seam cell divisions at the L3–L4 transition and epidermal fusion of seam cell daughters proceed largely normally, although final seam–seam fusion in L4 is incomplete in a subset of animals. **E.** Lineage schematics summarizing seam cell division patterns in control and seam cell–specific MYRF-1 AID animals. **F.** Reiteration of L2-like seam cell divisions induced by MYRF-1 AID is suppressed by loss of *hbl-1*. Quantification of seam cell numbers shows increased seam cell counts in MYRF-1 AID animals and suppression of this phenotype in *hbl-1* mutants.

We next examined expression of *dpy-4*, *dpy-5*, and *dpy-13*, L2-specific collagen genes^[41]^. In *myrf-1* single mutants, these genes were expressed and incorporated into the new cuticle, despite ecdysis failure (Figure 2A; Figure S2B). However, in *myrf-1/2* double mutants, expression of these collagens was absent in the anterior epidermis (hyp1–6) (Figure 2A; Figure S2B). A similar anterior defect was observed in *myrf-1(G274R)* mutants. A *dpy-5p* transcriptional reporter expressing TagRFP::PH confirmed loss of anterior epidermal promoter activity in *myrf-1/2* double mutants (Figure S2C), suggesting impaired regional patterning and transcriptional activation.

To assess broader aspects of cuticle remodeling, we examined K10D3.4, a serine protease inhibitor expressed in the pharynx and secreted into the pharyngeal cuticle during molt^[41]^ (Figure S2D). In *myrf-1* AID animals, K10D3.4 protein showed normal cycling between dispersion, secretion, and endocytosis. In contrast, MYRF-1 AID; *myrf-2(-)* animals displayed persistently diffuse K10D3.4, with no evidence of dynamic redistribution, and overall signal was significantly reduced (Figure S2D, E).

These data indicate that *myrf-1* single mutants initiate molting and synthesize L2 cuticle but fail to complete ecdysis, leading to cuticle retention. In contrast, *myrf-1/2* double mutants or MYRF-deficient animals display a more severe phenotype—aberrant or regionally incomplete L2 collagen gene expression and complete failure to initiate cuticle shedding. Together, these results demonstrate that MYRF activity is required both to drive the execution of molting and to ensure proper spatial patterning of the new cuticle during larval transitions (see section “MYRF Functions as an Amplifier of the Molting Oscillator”).

### Late-Stage-Specific Somatic Programs Require MYRF Beyond Molting

Seam cell development provides a well-established paradigm for larval stage identity in *C. elegans*, as these epidermal stem-like cells undergo a stereotyped sequence of stage-specific symmetric and asymmetric divisions that are tightly coupled to larval transitions^[42]^ (Figure 2B). Because each larval stage is marked by a distinct seam cell division pattern, deviations from this program serve as a sensitive and quantitative readout of defects in temporal progression^[7]^.

To assess how MYRF deficiency disrupts larval development, we examined stage-specific events across multiple somatic lineages, beginning with seam cells. In wild-type animals, seam cells V1–V4 and V6 undergo one asymmetric division in mid-L1, followed by a symmetric division and then a second asymmetric division around the L1–L2 molt, with additional asymmetric divisions at the L2–L3 and L3–L4 transitions. In *myrf-1* mutants or upon MYRF-1 depletion by AID, seam cells executed the late L1–early L2 divisions, although with altered timing (Figure 2B; Figure S2F, H). In contrast, in *myrf-1/2* double-deficient animals, these divisions failed to occur altogether, indicating a complete arrest of seam cell lineage progression prior to execution of the L2 program.

We next examined anchor cell (AC) and vulval development^[43–45]^, which initiate at L3 (Figure 2C; Figure S3A-D). The AC invades the basement membrane at the L2–L3 molt and fuses with the uterine seam cells (UTSE) in early L4. VPCs divide from late L3 through mid-L4, followed by vulval morphogenesis in late L4. Using *cdh-3p* and *zmp-1p* reporters, we observed that in MYRF-1 AID animals (induced from early L3), AC–UTSE fusion occurred, albeit with a delay, and P6.p divided and underwent toroid formation, indicating progression to early/mid-L4. In contrast, AC–UTSE fusion was never observed in MYRF-1 AID; *myrf-2(-)* animals, and P6.p failed to divide (Figure 2C; Figure S3C). Nevertheless, their gonads continued to develop, with oocyte-like cells observed—suggesting that germline expansion proceeds despite somatic arrest.

Upon MYRF-1 AID induction in early L4, we observed failed vulval eversion during late L4. This morphogenetic arrest was more pronounced in MYRF-1 AID; *myrf-2(-)* animals, in which morphogenesis did not initiate. *egl-17p::GFP*, marking vulF and vulE lineages from P6.p, confirmed vulval morphogenesis arrest (Figure S3B, D).

We also analyzed the M-cell lineage, which initiates from mid-L1 and gives rise to 16 descendants, including sex myoblasts (SMs)^[42]^. SMs migrate to the midbody in L2 and undergo stereotyped divisions during mid-to-late-L3 to form vulval muscle precursors. In all *myrf* mutants and MYRF-deficient animals, M-cell divisions and sex myoblast migration occurred with only modest delays; SMs completed their L3 divisions, indicating that M-lineage progression is comparatively less dependent on MYRF function (Figure S3E).

Taken together, these findings show that MYRF is essential for late-stage developmental events in epidermal and uterine lineages, whereas the germline and M-lineage can proceed independently, supporting a model of tissue-specific developmental uncoupling under MYRF deficiency.

### Seam Cell–Specific MYRF-1 Depletion Causes Heterochronic L2-Reiteration

Because global MYRF depletion causes seam cell division arrest, we next asked whether MYRF-1 acts cell-autonomously within the seam cell lineage. To address this, we used the AID system to degrade MYRF-1 specifically in seam cells (Figure 2D). Under seam cell specific MYRF-1 AID, seam cells progressed through the normal two rounds of divisions during the L1-L2 molt and early L2: first a symmetric division, followed by an asymmetric one. However, we observed an additional round of symmetric division in L3, which preceded the expected, one-round L3-specific asymmetric division, resulting in a significant increase in total seam cell number (Figure 2D; Figure S4A). Using an *hyp-*7p -expressed nuclear marker to track the fusion of differentiated seam cell daughters with the epidermis, we found that the following asymmetric divisions proceeded normally: the anterior daughter cells adopted epidermal fates and fused with hyp7 as expected, although the fusion was often delayed compared to controls (Figure S4B). The L4-specific asymmetric division proceeds normally in MYRF-1 AID animals; however, final seam cell fusion is incomplete in a subset of animals (Figure 2D; Figure S4C). These results indicate that MYRF-1 depletion in seam cells reiterates the L2-like division program during L3, reminiscent of a heterochronic phenotype (Figure 2E).

Notably, seam cell–specific MYRF-1 depletion also led to a highly penetrant vulval bursting phenotype (Figure S4D). A similar phenotype has been classically observed in *let-7* family mutants^[10]^. To explore the molecular basis of bursting in MYRF-1 AID animals, we performed RNAi knockdown of heterochronic pathway components and found that *hbl-1*, *lin-28*, and *lin-41* RNAi effectively suppressed the vulval bursting phenotype (Figure S4D). This is consistent with previous studies indicating that knockdown of these genes can suppress bursting in *let-7* mutants^[11]^.

Given that *hbl-1* encodes a transcription factor that specifies L2-stage cell fate in the heterochronic pathway^[8]^, we further tested whether *hbl-1* is genetically downstream of MYRF-1 in seam cell regulation. We found that an *hbl-1* loss-of-function mutation suppressed the excessive seam cell proliferation observed in MYRF-1 AID animals (Figure 2F; Figure S4E), indicating that *hbl-1* is a key downstream effector of MYRF-1 in controlling seam cell temporal progression.

### MYRF-1 Drives Activation of *let-7* Family Heterochronic microRNAs

Given the striking similarity between the seam cell phenotypes observed in seam cell–specific MYRF-1–depleted animals and *let-7* family microRNA (*mir-48 mir-241 mir-84*) triple mutants—most notably the reiteration of L2-like symmetric divisions^[8]^—and the fact that their primary target *hbl-1* functions downstream of MYRF-1 activity, we considered the possibility that MYRF-1 regulates the expression of these heterochronic microRNAs.

To explore this, we first analyzed the temporal relationship between MYRF-1 nuclear enrichment and microRNA expression (Figure 3). Previous work showed that *lin-4*, *mir-84*, and *let-7* transcripts display oscillatory expression across larval stages^[46]^, with peaks occurring shortly before molting—coinciding with the oscillatory peak of *myrf-1* mRNA expression. In contrast, *mir-48* and *mir-241* exhibit an earlier peak beginning in late L1 and further elevated in L2, then maintained at relatively stable levels thereafter^[46]^. Using endogenously tagged microRNA transcriptional reporter (with rapid degradation sequence) strains generated by the Caenorhabditis Genetics Center (CGC), we found that *mir-48p* and *mir-241p* reporters begin to increase in expression coinciding with MYRF-1 nuclear entry during late L1. *Let-7p* reporter expression was also clearly detectable in late L1 (Figure 3A). In contrast, *mir-84p* reporter expression remained minimal in Late L1 and showed a strong upregulation in late L2, also aligned with the nuclear peak of MYRF-1.

**Figure 3.**
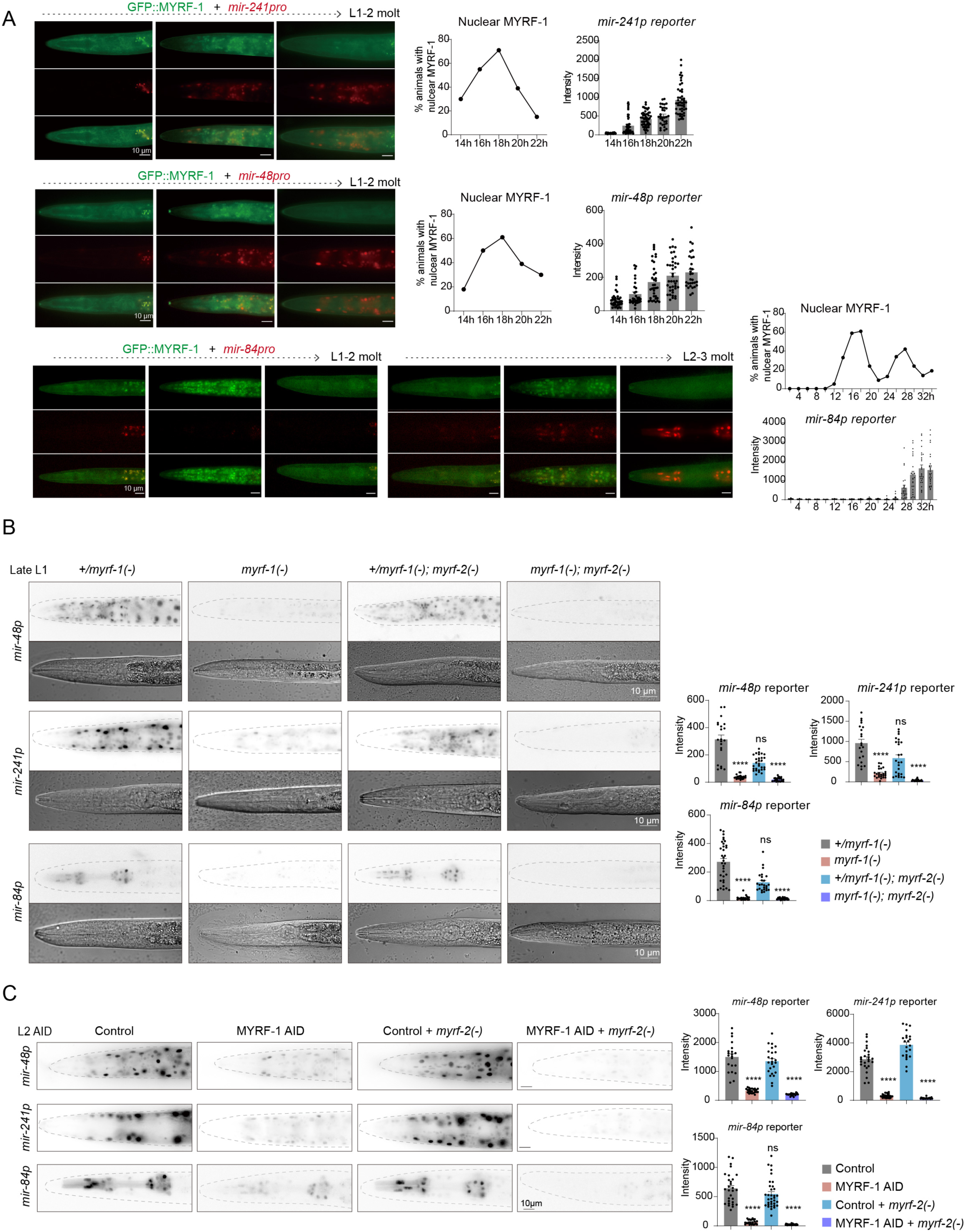
MYRF drives temporal activation of *let-7* family heterochronic microRNAs. **A.** Imaging of GFP::MYRF-1 (green) and transcriptional reporters for *mir-241*, *mir-48*, and *mir-84* (red) across the L1–L2 and L2–L3 transitions. Nuclear accumulation of MYRF-1 peaks during late intermolt phases and precedes with activation of *mir-241* and *mir-48* reporters in late L1, whereas *mir-84* reporter activation occurs predominantly in late L2. Quantification shows the fraction of animals with nuclear MYRF-1 and corresponding reporter intensities over time. **B.** Endogenous *mir-48*, *mir-241*, and *mir-84* reporter expression in late L1 animals of indicated genotypes. Reporter activity is strongly reduced in *myrf-1* mutants and abolished in *myrf-1; myrf-2* double mutants. Right, quantification of reporter intensities. **C.** Requirement for MYRF-1 during late L2. MYRF-1 AID initiated in early L2 markedly suppresses *mir-48*, *mir-241*, and *mir-84* reporter expression, even after initial induction has begun. Quantification is shown on the right.

To test the requirement for MYRF-1, we analyzed microRNA reporter activity in *myrf-1* and *myrf-2* single mutants, *myrf-1/2* double mutants, and in animals subjected to MYRF-1 depletion from early L1 with or without *myrf-2(-)* (Figure 3B; Figure S5A, B). Loss of *myrf-1* alone caused a marked reduction in *mir-48p*, *mir-241p*, and *mir-84p* reporter activity, whereas expression of all three reporters was completely abolished in *myrf-1/2* double-deficient animals. *let-7* reporter expression was also significantly reduced upon MYRF-1 depletion (Figure S5C); however, due to technical limitations, its status could not be assessed in the *myrf-2(-)* background. Consistent with our previous findings that *lin-4* expression was eliminated in *myrf-1/2* double mutants, we confirmed that its reporter remained silenced in MYRF-deficient animals (Figure S5D).

Because *mir-48/241/84* reporters reach maximal expression during late L2 according to systematic RNA-seq profiles, we next asked whether MYRF-1 activity is required during this later window. We therefore initiated MYRF-1 degradation from early L2, after *mir-48/241/84* induction had already begun. Even under these conditions, reporter activity for *mir-48*, *mir-241*, and *mir-84* was strongly suppressed (Figure 3C), indicating that MYRF-1 activity during late L2 is required not only to initiate but also to maintain or amplify expression of these microRNAs.

Taken together, these results support a model in which MYRF-1 promotes the timely activation and reinforcement of *let-7* family microRNAs—including *mir-48*, *mir-241*, and *mir-84*—as well as *let-7* itself. Failure to activate these microRNAs in seam cell–specific MYRF-1–deficient animals likely underlies the reiteration of L2-specific seam cell divisions, supporting a cell-autonomous role for MYRF-1.

### MYRF-1 Directly Targets on Heterochronic miRNA and Developmental Regulators

To identify the direct transcriptional targets of MYRF-1, including microRNA genes, we collaborated with the modERN project^[47]^ to perform MYRF-1 ChIP–seq analysis (Figure 4). Synchronized late L2 larvae (27 h post-hatching) were collected for genome-wide profiling. This analysis revealed that the majority of MYRF-1 binding sites are located within ∼1 kb upstream of transcription start sites (TSSs), consistent with MYRF-1 functioning primarily as a promoter-proximal transcriptional regulator (Figure 4A-C).

**Figure 4.**
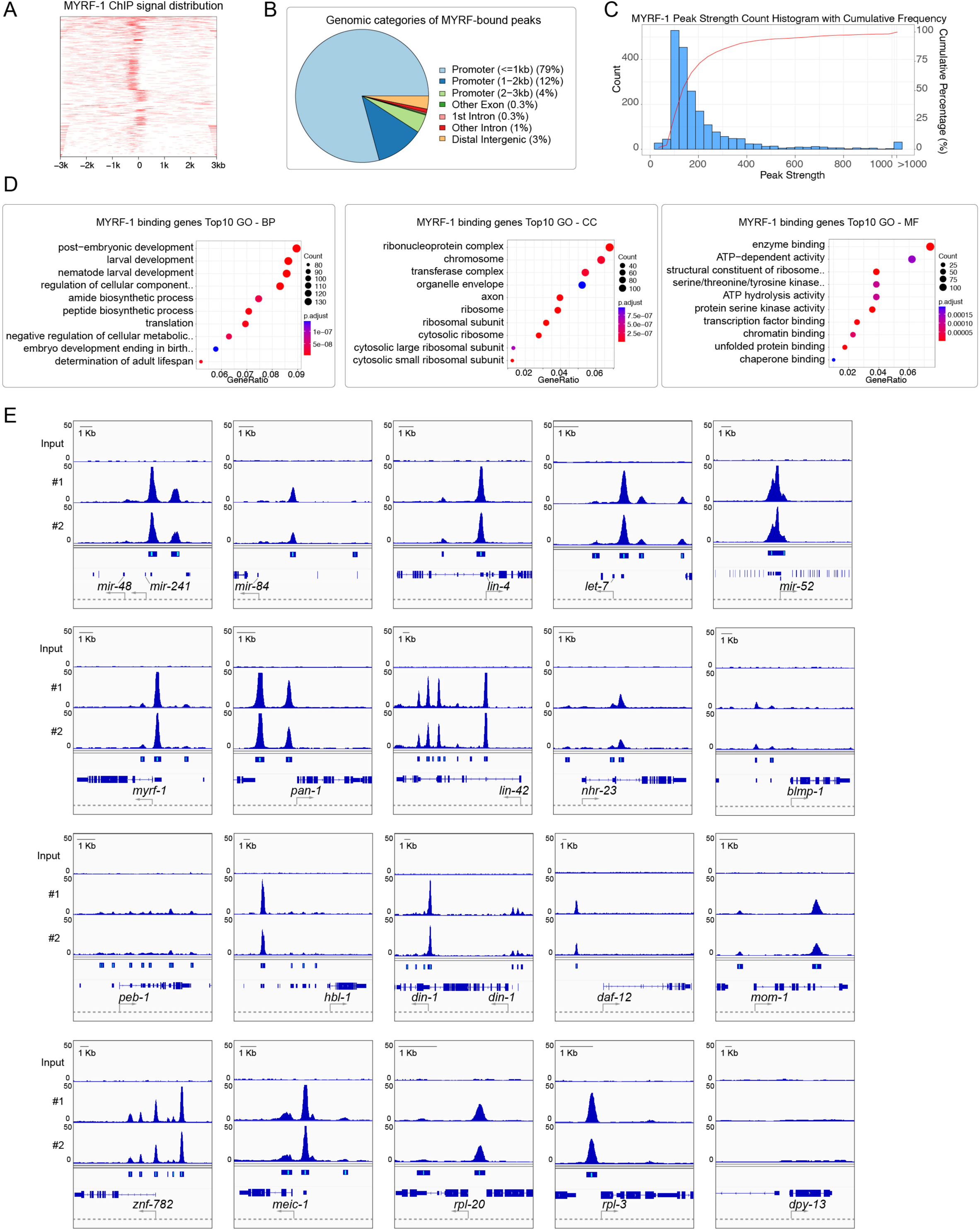
Genome-wide identification of MYRF-1 direct transcriptional targets by ChIP–seq. **A.** Heatmap of MYRF-1 ChIP–seq signal distribution showing strong enrichment at promoter-proximal regions. **B.** Genomic distribution of MYRF-1–bound peaks. The majority of binding sites localize within promoter regions (<1 kb upstream of transcription start sites), consistent with a role as a promoter-proximal transcriptional regulator. **C.** Histogram of MYRF-1 peak strengths with cumulative frequency, indicating a small number of high-occupancy peaks and a broader distribution of moderate binding sites. **D.** Gene Ontology (GO) enrichment analysis of MYRF-1–bound genes. Top enriched terms include biological processes related to post-embryonic and larval development (BP), ribonucleoprotein and ribosomal complexes (CC), and transcriptional and enzymatic regulatory activities (MF). **E.** Genome browser views of representative MYRF-1 ChIP–seq peaks at heterochronic microRNA loci (*mir-48/241*, *mir-84*, *lin-4*, *let-7*), oscillatory timing regulators (*lin-42*, *nhr-23*, *blmp-1*), and additional developmental or translational regulatory genes. In contrast, no MYRF-1 binding is detected at cuticle structural genes (e.g., *dpy-13*). Two independent ChIP replicates (#1, #2) and input controls are shown.

Notably, strong MYRF-1 binding peaks were detected at the loci of key heterochronic microRNAs, including the *mir-48/241*, *mir-84*, *let-7*, and *lin-4*, among the most prominent peaks in the dataset, strongly suggesting direct transcriptional regulation (Figure 4D, E). In addition, robust MYRF-1 occupancy was observed near the embryonically and early larval–expressed miRNAs *mir-52* and the *mir-54/55/56* family (Figure 4E) (File S1), suggesting that MYRF-1 may also regulate additional miRNA modules active during early developmental stages.

Beyond miRNAs, MYRF-1 binding sites were mapped to approximately 2,400 protein-coding genes (File S2). Gene Ontology (GO) enrichment analysis revealed a highly significant overrepresentation of genes involved in larval development^[48]^, encompassing more than 130 genes (File S3). This group includes classical heterochronic regulators (e.g. *lin-14*, *lin-41*, *lin-28*, *hbl-1*, *lin-42*, *daf-12*); core molting cycle regulators (e.g. *nhr-23*, *blmp-1*, *grh-1*, *peb-1*); Wnt signaling components (e.g. *mom-1*, *lit-1*, *wrm-1*); transcriptional regulators and chromatin modifiers (e.g. *egl-27*, *lin-9*, *lin-35/Rb*, *set-16*, *sop-2*); major signaling pathway components (e.g. *mpk-1*, *let-60/Ras*, *pkc-1*, *kin-29*); genes involved in neuronal development and synaptic function (e.g. *unc-108*, *unc-104*, *unc-32*, *egl-3*); and regulators of nutritional and metabolic state (e.g. *daf-16*, *aak-2*, *daf-2, daf-5*, *din-1*). Together, these targets illustrate an potential integration of transcriptional regulation, signaling pathways, molting machinery, and metabolic state in orchestrating larval development. Notably, strong MYRF-1 binding was also detected at the *myrf-1*, *myrf-2*, and *pan-1* loci (Figure 4E), suggesting the existence of autoregulatory and feedback control within the MYRF regulatory network.

In addition to developmental genes, MYRF-1 targets were significantly enriched for cytosolic ribosome components (Figure 4D, E) (File S3), including numerous ribosomal protein genes (e.g. *rpl-3*, *rpl-5*, *rps-3*, *rps-10*) and translational regulators (e.g. *gld-4*, *cgh-1*, *pab-2*, *sup-26*, *eIF-3* complex), suggesting that MYRF-1 also engages the general translational machinery.

### MYRF Functions as an Amplifier of the Molting Oscillator

To characterize the transcriptional landscape regulated by MYRF, we performed mRNA profiling of MYRF-1 AID; *myrf-2(−)* animals initiated from early L1, with untreated animals as controls. Samples were collected at a late L1 stage corresponding to the nuclear peak of MYRF-1 in control animals, a time window expected to capture key MYRF-dependent regulatory events.

RNA-seq analysis identified 1,281 downregulated genes and 691 upregulated genes upon MYRF depletion (Figure 5A) (File S4). The most significantly enriched GO category among downregulated genes was molting cycle^[48]^ (Figure 5; Figure S6A) (File S5), including cuticle collagen genes (e.g. *dpy-3, dpy-4, dpy-5, dpy-13*), precuticle components (e.g. *noah-1, noah-2*), proteases and remodeling enzymes (e.g. *nas-36, nas-37, nas-38, bli-5*), PTR/PATCHED-related transmembrane proteins involved in cuticle patterning (e.g. *ptr-1, ptr-2, ptr-4, ptr-9*), and core molting regulators (e.g. *nhr-23, grh-1, bed-3, myrf-1, pan-1, blmp-1, dre-1*). Notably, the downregulation of *myrf-1* and *pan-1* (Figure 5E), together with strong MYRF-1 occupancy at the *myrf-1* and *pan-1* loci, indicates a positive self-reinforcing feedback mechanism that elevates the oscillatory peak of *myrf-1* expression. This broad suppression of molting-associated genes upon MYRF degradation is consistent with the molting defects and larval arrest observed in *myrf*-deficient animals, highlighting a crucial role for MYRF in coordinating the temporal activation of the molting program. In contrast, prominently upregulated genes included members of the cytochrome P450 family (e.g. *cyp-33 and cyp-34* subfamilies) and glutathione S-transferases (e.g. *gst-14, gst-15*) (Figure 5B, E), enzymes broadly associated with lipid metabolism and detoxification.

**Figure 5.**
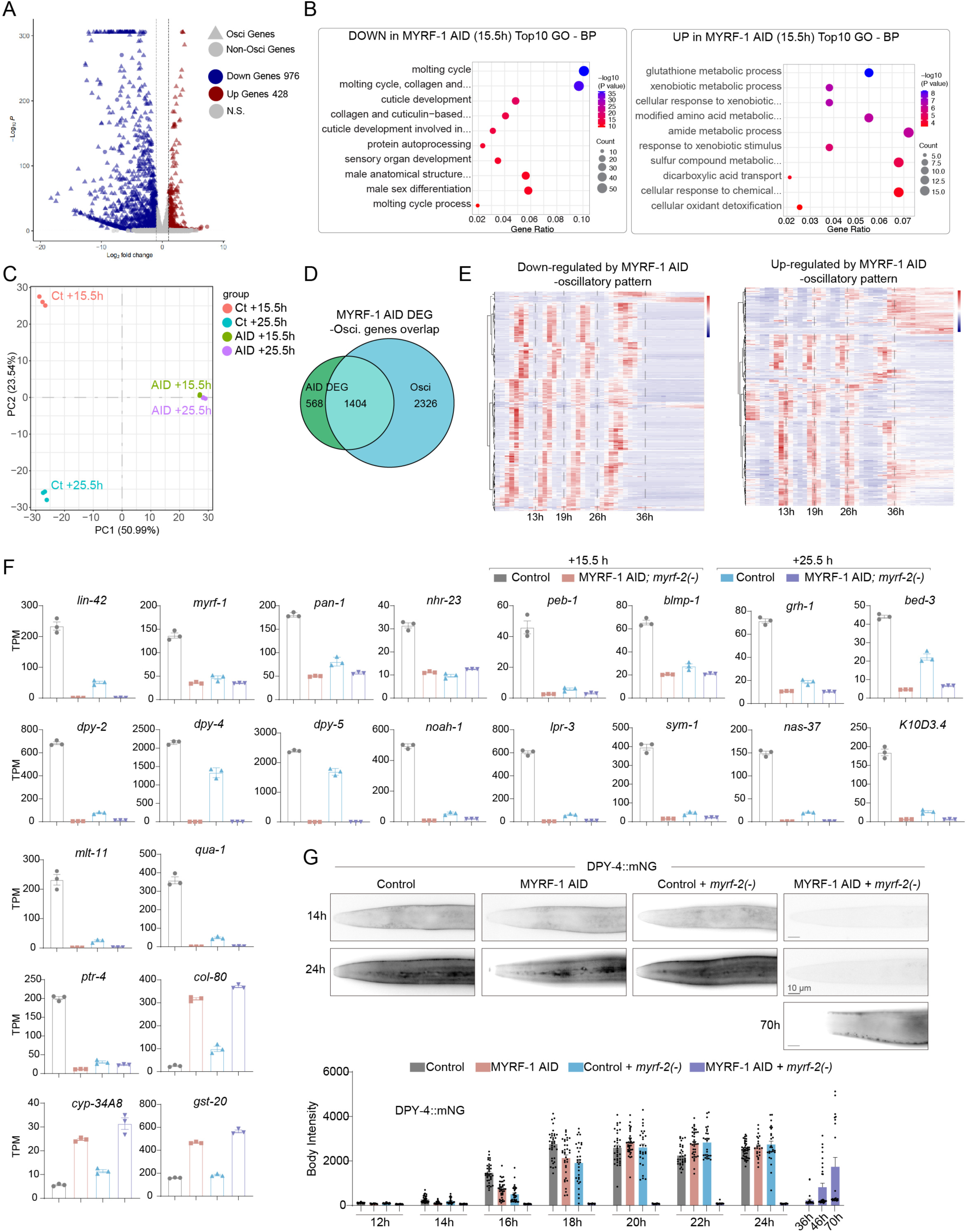
MYRF is required to promote the oscillatory gene network and molting program. **A.** Volcano plot showing differentially expressed genes following MYRF-1 AID at 15.5 h. Oscillatory (Osci) and non-oscillatory genes are highlighted. Blue, downregulated genes; red, upregulated genes; gray, not significant. **B.** Gene Ontology (GO) enrichment analysis of genes downregulated (left) or upregulated (right) upon MYRF-1 AID at 15.5 h. BP (Biological Processes) of GO is shown. Downregulated genes are strongly enriched for molting cycle, cuticle organization, and larval development, whereas upregulated genes are enriched for detoxification and metabolic processes. **C.** Principal component analysis (PCA) of RNA-seq samples from control and MYRF-1 AID animals at 15.5 h and 25.5 h. MYRF-1 AID samples cluster closely at both time points, indicating high similarity, whereas control samples are clearly separated, reflecting normal temporal progression. **D.** Overlap between MYRF-1 AID differentially expressed genes (DEGs) and previously defined oscillatory genes by Meeuse *et al.*, indicating preferential disruption of the oscillatory transcriptome. **E.** Heatmaps showing temporal expression patterns of oscillatory genes downregulated (left) or upregulated (right) by MYRF-1 AID across larval development, revealing distinct phase relationships between the two gene groups. Temporal expression profiles are based on datasets from Meeuse *et al*. **F.** RNA-seq quantification of representative MYRF-dependent genes. Oscillatory regulators (*lin-42, nhr-23, peb-1, blmp-1, grh-1, bed-3*), molting and cuticle genes (*dpy-2, dpy-4, dpy-5, noah-1, lpr-3, mlt-11, nas-37*), and detoxification genes (*gst-20, cyp-34A8, col-80*) are shown at 15.5 h and 25.5 h in control, MYRF-1 AID, and MYRF-deficient backgrounds. **G.** DPY-4::mNG reporter analysis following MYRF depletion. Representative images (top) and quantitative fluorescence measurements (bottom) show robust DPY-4 induction in control animals but failure of normal activation in MYRF-deficient animals. In arrested animals, DPY-4 protein accumulates slowly over time but remains absent from the anterior epidermis (head region).

Importantly, downregulated structural cuticle genes generally lack direct MYRF-1 binding sites, indicating that the effect is largely indirect. Instead, MYRF-1 directly targets key transcriptional regulators of the molting program, including *nhr-23*, *blmp-1*, *grh-1*, and *peb-1*, among which *nhr-23* displays particularly strong MYRF-1 occupancy (Figure 5E; Figure 4E). This suggested that MYRF may function upstream to promote the molting transcriptional cascade.

To examine this relationship, we generated an endogenous *nhr-23::mKate2* reporter and compared its dynamics with GFP::MYRF-1 (Figure 6A). Nuclear accumulation of NHR-23 preceded MYRF-1 nuclear enrichment by a short interval. NHR-23 expression was largely restricted to epidermal cells, whereas MYRF-1 was broadly expressed. At the onset of NHR-23 expression, MYRF-1 remains predominantly membrane-localized across tissues. Subsequently, MYRF-1 began to accumulate in epidermal and seam cell nuclei while retaining partial membrane localization, before transitioning to a predominantly nuclear distribution. The decline of NHR-23 expression also slightly preceded the reduction of nuclear MYRF-1 levels.

**Figure 6.**
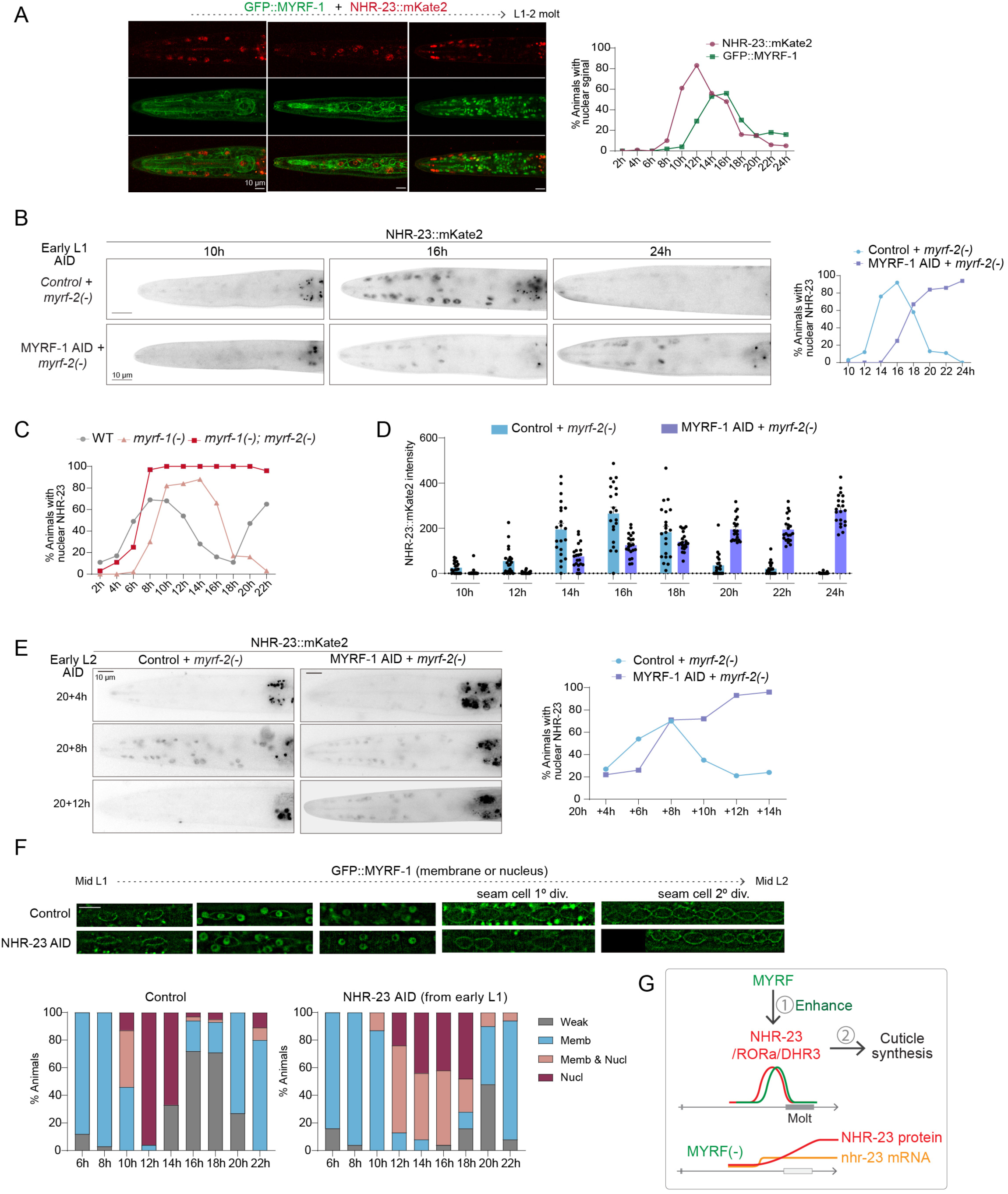
MYRF amplifies NHR-23 expression to drive molting oscillations. **A.** Imaging of GFP::MYRF-1 (green) and NHR-23::mKate2 (red) during the L1–L2 transition. NHR-23 nuclear accumulation precedes that of MYRF-1. MYRF-1 nuclear localization first appears in seam and epidermal cells while membrane-associated MYRF-1 remains detectable, and subsequently becomes broadly nuclear with minimal membrane signal. Right, quantification of the fraction of animals exhibiting nuclear MYRF-1 or NHR-23 over time. **B.** NHR-23::mKate2 expression following early-L1 MYRF-1 AID in a *myrf-2(–)* background. Representative images and quantification (right) of the fraction of animals with nuclear NHR-23 during the L1–L2 interval show delayed yet sustained nuclear accumulation of NHR-23 compared with controls. **C.** Quantification of the fraction of animals with nuclear NHR-23 in WT, *myrf-1(–)*, and *myrf-1(–); myrf-2(–)* animals. Oscillatory nuclear accumulation is retained in *myrf-1(–)* mutants but is flattened in *myrf-1/2* double mutants. **D.** Quantification of NHR-23::mKate2 fluorescence intensity over time in control and MYRF-deficient animals, revealing delayed onset and altered dynamics of NHR-23 accumulation. **E.** NHR-23::mKate2 dynamics following early-L2 MYRF-1 AID. Representative images and quantification indicate delayed, prolonged NHR-23 nuclear accumulation and loss of oscillatory decline. **F.** GFP::MYRF-1 localization relative to seam cell divisions in control and NHR-23 AID animals^[28]^. Quantification below shows prolonged nuclear MYRF-1 residence upon NHR-23 depletion. **G.** Model illustrating MYRF function as an amplitude amplifier of NHR-23 transcription. MYRF enhances NHR-23 expression to drive cuticle synthesis and molting progression; in MYRF-deficient animals, reduced transcriptional amplitude results in sustained but ineffective NHR-23 protein accumulation, leading to oscillatory arrest.

In MYRF-1 AID; *myrf-2(-)* from early L1, NHR-23::mKate2 signals were not abolished but delayed: it required approximately 5–7 hours to reach levels comparable to controls (Figure 6B). Strikingly, the normal oscillatory decline of NHR-23 was not observed; instead, NHR-23 signals persisted in arrested animals (Figure 6B, D). A delayed and plateaued NHR-23 expression was also observed in *myrf-1/2* double mutants (Figure 6C). In *myrf-1* single mutants, NHR-23 expression was delayed but still exhibited a decline, albeit with a broadened peak window. When MYRF depletion was initiated from early L2, a comparable delay and persistence of NHR-23 was observed in late L2 (Figure 6E), consistent with *myrf-1* and *nhr-23*’s oscillatory expression patterns across larval cycles. Together, these data indicate that MYRF is not required to initiate *nhr-23* expression, but is essential for boosting its amplitude and ensuring proper oscillatory dynamics.

Consistent with imaging results, RNA-seq analysis of MYRF-deficient animals at late L1 (15.5 h) revealed that *nhr-23* transcripts were reduced to ∼30% of control levels (Figure 5E), a stage at which NHR-23::mKate2 peaked in controls but was barely detectable in MYRF-depleted animals. Because NHR-23::mKate2 levels later increased in MYRF-deficient animals, we reasoned that the suppressed molting program observed at 15.5 h might be corrected after additional hours of development through delayed accumulation of NHR-23 protein. We therefore performed a second RNA-seq analysis at 25.5 h (Figure 5C, E), when NHR-23::mKate2 levels in MYRF-depleted animals approached those of controls at 15.5 h. Surprisingly, the 25.5 h RNA-seq did not reveal increased *nhr-23* mRNA or recovery of molting gene transcripts (Figure 5E). Principal component analysis (PCA) showed that the two MYRF-deficient transcriptomes clustered closely (Figure 5C), indicating minimal global transcriptional change between these time points, whereas control animals exhibited pronounced transcriptomic shifts consistent with normal progression from late L1 to mid-L2. Together, these results suggest that NHR-23 protein accumulation in arrested animals reflects slow protein buildup from persistently low transcription rather than restoration of normal transcriptional output (Figure 6G).

Consistent with this interpretation, reporters such as *dpy-4::mNG* and *dpy-5::mNG* were eventually detected in MYRF-deficient animals despite low transcript levels by RNA-seq (Figure 5E). GFP signals were absent at the time corresponding to peak expression in controls but gradually accumulated after prolonged arrest (>24 h) (Figure 5F; Figure S6B). Together, these results support a model in which MYRF is not required for molting initiation but is essential for amplifying the molting transcriptional program.

Because molting-related genes constitute a major subset of the larval oscillatory transcriptome, we examined the overlap between oscillatory genes and differentially expressed genes (DEGs) following MYRF depletion at 15.5 h. Strikingly, more than 70% of MYRF-dependent DEGs (1404 of 1972) correspond to previously defined oscillatory genes (Figure 5D) (File S6). These genes include both downregulated and upregulated groups upon MYRF-1 AID. When their temporal expression profiles were clustered using the published oscillation dataset from Meeuse *et al.*^[24]^, a clear phase-specific pattern emerged: genes downregulated by MYRF-1 AID normally peak during the mid-to-late intermolt period, whereas genes upregulated upon MYRF depletion peak either earlier or later in the oscillatory cycle (Figure 5E). These results indicate that MYRF broadly reshapes the oscillatory gene expression landscape in a phase-selective manner, preferentially amplifying transcriptional programs associated with late-intermolt progression.

### MYRF-1 Activates *lin-42* and Is Restrained by *lin-42* Feedback

*lin-42*, the *C. elegans* homolog of the circadian Period gene, exhibits oscillatory expression that peaks immediately prior to molting in each cycle. Loss of *lin-42* disrupts the synchrony of molting phases, and *lin-42* is therefore considered a core component of the larval developmental oscillator. Our ChIP-seq analysis revealed that the *lin-42* locus displays the strongest MYRF-1 occupancy among all identified targets (Figure 4E) (File S2). Consistently, RNA-seq showed that *lin-42* transcripts are drastically reduced in MYRF-deficient animals (Figure 5E) (File S4), prompting us to examine in detail how MYRF regulates *lin-42* expression.

We generated an endogenous *lin-42::mKate2* knock-in strain and analyzed its expression dynamics alongside GFP::MYRF-1 (Figure 6A). We found that the peak of nuclear MYRF-1 accumulation consistently precedes the emergence of *lin-42* nuclear signal by approximately two hours. Following MYRF-1 downregulation during lethargus, *lin-42* expression also declined. Both MYRF-1 and *lin-42* exhibited broad tissue expression, including epidermal cells, pharynx, body wall muscles, and neurons.

In *myrf-1* null animals, LIN-42::mKate2 expression was markedly reduced and its peak delayed, indicating impaired activation during late L1 (Figure 7B). In *myrf-1/2* double mutants, LIN-42 expression was completely abolished at late L1, demonstrating that MYRF proteins are essential for *lin-42* activation. When MYRF depletion was initiated in early L2, LIN-42 protein levels declined from the late L1 peak in both control and AID-treated animals, and then rose again at late L2 in controls. In contrast, LIN-42 expression remained suppressed under MYRF-1 AID conditions (Figure S7A), indicating that MYRF is required not only for initial *lin-42* activation but also for its oscillatory reactivation during subsequent larval stages.

**Figure 7.**
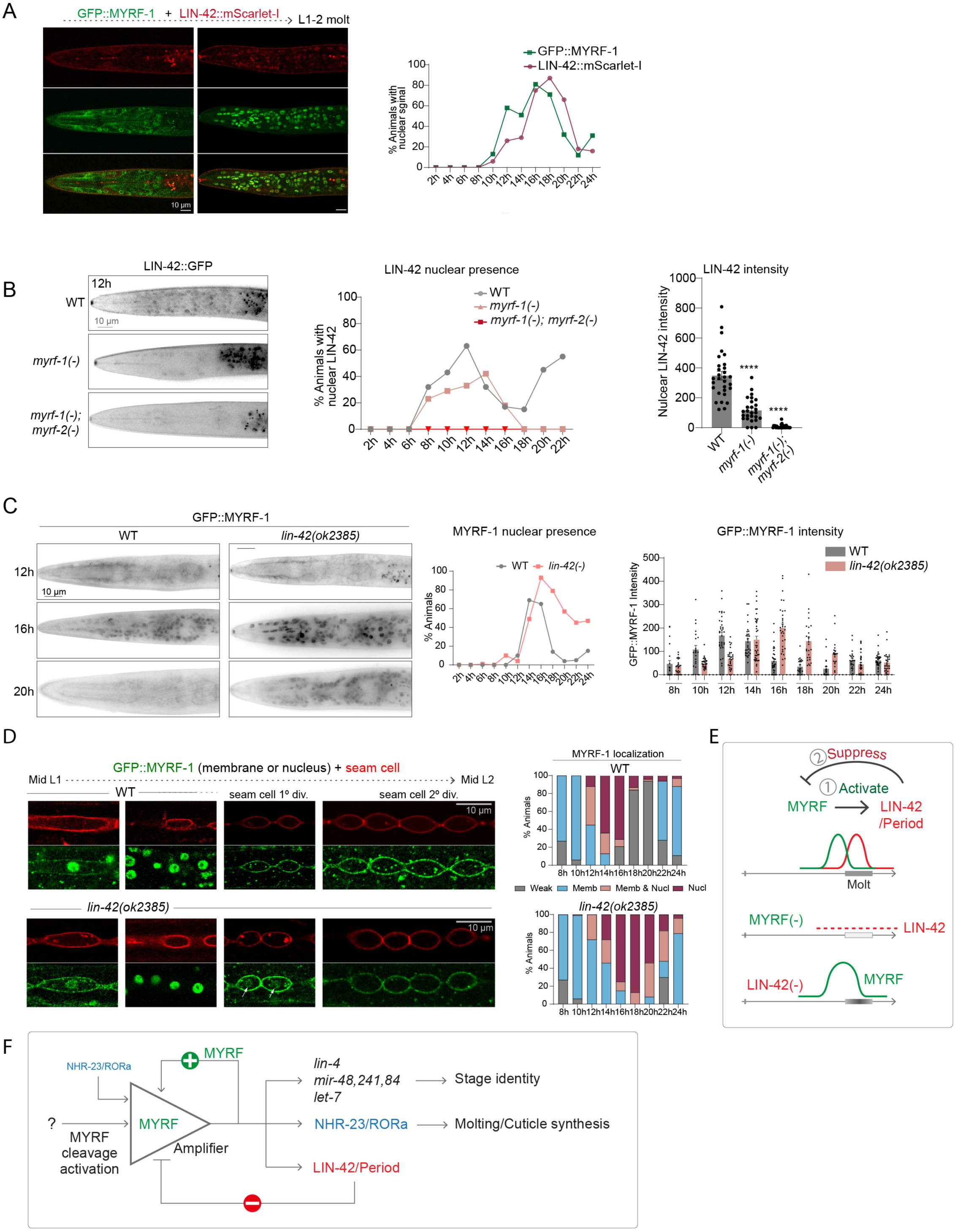
MYRF activates LIN-42 to sharpen oscillatory phases. **A.** Imaging of GFP::MYRF-1 (green) and LIN-42::mScarlet-I (red) during the L1–L2 transition. LIN-42 nuclear accumulation follows the peak of nuclear MYRF-1. Right, quantification of the fraction of animals with nuclear MYRF-1 or LIN-42 over time. **B.** LIN-42::GFP expression in WT, *myrf-1(–)*, and *myrf-1(–); myrf-2(–)* animals. Left, representative images at 12 h. Middle, fraction of animals with nuclear LIN-42 over time. Right, quantification of nuclear LIN-42 intensity, showing reduced activation in MYRF-deficient backgrounds. **C.** GFP::MYRF-1 localization in WT and *lin-42(–)* animals. Left, representative images at indicated times. Middle, fraction of animals with nuclear MYRF-1 over time. Right, quantification of GFP::MYRF-1 intensity, revealing prolonged nuclear residence in *lin-42* mutants. **D.** Seam cell–focused imaging of GFP::MYRF-1 localization (green) with seam cell membrane markers (red) from mid-L1 to mid-L2. In *lin-42(–)* animals, nuclear MYRF-1 persists in seam cells during the first round of symmetric division (arrow), whereas nuclear MYRF-1 is rarely observed at this stage in controls. Quantification (right) shows prolonged nuclear MYRF-1 localization in *lin-42(–)* animals compared with controls. **E.** Model of MYRF–LIN-42 regulation. MYRF activates *lin-42/Period*, which in turn suppresses *myrf* with a phase delay, sharpening oscillatory dynamics during the molting cycle. Loss of either component disrupts oscillatory program. **F.** Model of MYRF-centered control of larval stage transitions. Activated MYRF functions as an amplifier with positive self-feedback, coordinating larval development by activating heterochronic microRNAs for stage identity and enhancing NHR-23/RORα for molting and cuticle synthesis. MYRF also induces LIN-42/Period, which feeds back negatively to constrain MYRF activity, forming a gated oscillatory circuit that couples stage identity with molting execution.

We next asked whether LIN-42, in turn, influences the oscillatory behavior of MYRF-1. In *lin-42(ok2385)* strong loss-of-function mutants^[20]^, we observed a pronounced extension of the MYRF-1 nuclear phase (Figure 7C). Although this could, in principle, reflect a general developmental delay, closer examination of seam cell behavior argued against this interpretation. In wild-type animals, during the first round, symmetric seam cell division at the L1–L2 transition, MYRF-1 is typically membrane-associated and never nuclear (Figure 7D). In contrast, in *lin-42* mutants, nuclear MYRF-1 signals were frequently observed during the symmetric seam cell division, indicating prolonged nuclear residence of MYRF-1 that is uncoupled from normal developmental progression. However, this extended nuclear localization was not maintained during the subsequent asymmetric division (Figure 7D), suggesting that LIN-42 contributes to MYRF-1 oscillatory dynamics in a stage-restricted and partial manner.

To test whether *lin-42* is sufficient to accelerate MYRF-1 degradation, we overexpressed *lin-42* in the intestine; however, this manipulation did not lead to a detectable reduction in MYRF-1 nuclear signal (Figure S7B). This suggests that LIN-42-mediated regulation of MYRF-1 likely operates through indirect or developmentally coupled mechanisms, rather than via direct degradation.

The transiently sustained nuclear MYRF-1 pattern observed in *lin-42* mutants contrasts with MYRF-1 localization in animals subjected to NHR-23 depletion. In NHR-23 AID animals, MYRF-1 nuclear presence is also prolonged (Figure 6F), but the decline of nuclear MYRF-1 always precedes symmetric seam cell division, suggesting with an overall developmental delay or arrest.

Taken together, these observations place MYRF-1 upstream of *lin-42* in the developmental timing hierarchy (Figure 7E). Given the central role of *lin-42* in coordinating oscillatory timing and molting physiology, its failure to be properly activated in MYRF-deficient animals likely contributes to the developmental arrest and heterochronic phenotypes observed.

## Discussion

Here, we identify MYRF-1 as a key regulator that integrates multiple temporal control systems governing larval development in *C. elegans* (Figure 7F). Our results show that MYRF-1 promotes heterochronic microRNA activation and functions as a self-reinforcing amplifier of oscillatory transcriptional programs that drive execution of the molting cycle and orderly stage transitions. Loss of MYRF disrupts each of these processes, leading to predictable, stage-specific developmental arrests. Together, these findings position MYRF-1 as a central hub coordinating larval timing and stage progression.

A major finding of this work is that MYRF-1 is required to activate the expression of the heterochronic microRNAs. Analysis using endogenous expression reporters reveals that *lin-4*, *mir-48/241,* and *let-7* are strongly induced in late L1, whereas *mir-84* is predominantly activated in late L2. Once activated, transcription from these microRNA loci persist and exhibit oscillatory expression across all four larval stages. Inactivation of *myrf* not only abolishes the initial activation of these microRNAs but also suppresses their expression in later stages. Consistent with this loss, *myrf* deficiency leads to characteristic heterochronic phenotypes, including sustained *lin-14* expression (associated with loss of *lin-4*), reiteration of L2-like seam cell divisions (associated with loss of *mir-48/241/84*), and vulval bursting (associated with loss of *let-7*). Given the strong binding of MYRF at these microRNA loci—and the relative scarcity of comparably strong binding at other genes—activation of heterochronic microRNAs appears to be a central function of MYRF in driving stage identity transitions.

Notably, mature *mir-48/241* levels do not peak until L2–L3, coinciding with their functional roles in regulating L3-stage events. Similarly, although *let-7* is also transcriptionally activated in late L1, its functional activity is delayed until L4. This temporal separation between transcription and functional output suggests additional regulatory layers governing *mir-48/241* activity. Early transcription may serve to preload or buffer microRNAs before their concentrations exceed the threshold required for effective target repression, or *mir-48/241* maturation may require additional post-transcriptional regulation, analogous to LIN-28–mediated inhibition of *let-7*.

A second major conclusion is that MYRF-1 is essential for progression of the oscillatory transcriptional program during the late intermolt phase, including genes associated with molting. Importantly, MYRF-1 does not bind structural components of the cuticle or molting machinery directly. Instead, MYRF-1 binds loci encoding transcription factors such as *nhr-23*, *blmp-1*, and *grh-1*, which themselves exhibit strong oscillatory expression and are functionally critical for molting. In the case of *nhr-23*, MYRF-1 is not required to initiate transcription but is essential for boosting expression amplitude. In MYRF-deficient animals, *nhr-23* transcripts persist at a moderate basal level, while the protein slowly accumulates to a relatively high and sustained level over time. We propose that this repressed, flattened transcriptional profile fails to reach the amplitude threshold required to trigger downstream transcriptional waves or engage feedforward mechanisms, thereby halting oscillatory progression. Thus, MYRF-1 acts as a transcriptional amplifier that elevates oscillatory regulators beyond a critical threshold necessary for robust stage transitions.

Our data further reveal that MYRF-1 directly binds the *lin-42* locus with high occupancy and is required for its activation. *lin-42*, the homolog of the circadian clock gene *Period*, functions as a temporal brake that defines phase length, particularly during the late intermolt period. Activation of *lin-42* subsequently represses *myrf* with a phase delay, forming a negative feedback loop. Analysis of MYRF-1 membrane-to-nuclear dynamics relative to seam cell divisions indicates that the decline of nuclear MYRF is delayed relative to cellular progression in the absence of *lin-42*, revealing partial decoupling of MYRF oscillation from developmental timing. However, this delay is not sustained, and overexpression of *lin-42* alone does not markedly suppress nuclear MYRF, indicating that additional mechanisms contribute to MYRF oscillatory downregulation. Nevertheless, by activating *lin-42*, MYRF participates in a feedback-regulated developmental pacemaker that helps sharpen temporal gene expression waves.

The MYRF-dependent control of oscillatory regulators provides a mechanistic basis for the stage-specific arrest phenotypes seen in *myrf-1/2*–deficient animals. Epidermal molting exemplifies how MYRF, despite its ubiquitous expression, drives tissue-specific developmental programs by acting through epidermal regulators such as *nhr-23*. Consistent with this model, multiple late larval developmental events—including DD synaptic remodeling, seam cell divisions, anchor cell–uterine interactions, and vulval morphogenesis—fail to progress in the absence of MYRF. The precise downstream effectors mediating these arrests in each tissue remain to be defined. These late stage–specific cellular progression programs, together with activation of heterochronic microRNAs, constitute key outputs of MYRF activity and position MYRF as a core integrator of stage identity, molting cycle execution, and tissue-specific differentiation during larval stage transitions.

Notably, not all developmental programs are equally affected by MYRF deficiency. For example, the M-cell lineage progresses relatively normally, and the germline can continue development, indicating the existence of partially independent or parallel timing programs.

A remaining question is what mechanism activates MYRF cleavage. In parallel work, we have shown that MYRF cleavage is tightly constrained by dual inhibitory mechanisms that restrict its activation to defined developmental windows (Xu Z. et al., manuscript submitted)^[49]^. Disruption of these controls leads to premature activation of developmental programs, including early *lin-4* expression, and bypass of starvation-induced developmental arrest. These findings establish MYRF cleavage as a gatekeeper of larval stage transitions. However, the upstream positive signal that triggers MYRF cleavage remains elusive. Given MYRF’s broad expression across somatic tissues and the largely synchronous timing of its cleavage, this activating cue is likely systemic.

Such a systemic cue is likely linked to overall growth status, including germline expansion. A key observation from this study is that MYRF deficiency produces an “uncoupled growth arrest” during late intermolt, in which somatic developmental programs stall while body growth continues. This indicates that growth itself is not directly controlled by MYRF. Instead, under normal conditions, growth state likely provides instructive signals that promote progression of developmental programs^[50,51]^. We propose that MYRF functions as a central conduit for this communication: disruption of MYRF activity decouples developmental progression from growth. Conversely, although *myrf*-deficient animals initially continue to grow, growth ultimately ceases, indicating that sustained growth also depends on ongoing developmental progression. Together, these findings point to a bidirectional coupling between growth and developmental timing, with MYRF positioned at the interface of these processes.

## Acknowledgement

We thank Jordan Ward and Andrew Chisholm and for generously sharing collagen knock-in reporter strains and for insightful discussions that were instrumental in defining and interpreting molting phenotypes. We thank the Caenorhabditis Genetics Center (CGC) for providing endogenous microRNA reporter strains used in this study. We thank Michelle Kudron and Robert Waterston for performing MYRF-1 ChIP experiments through the modERN project, which yielded informative insights. We thank the Molecular and Cell Biology Core Facility (MCBCF) and the Molecular Imaging Core Facility (MICF) at the School of Life Science and Technology, ShanghaiTech University for providing technical support. Some strains were provided by the CGC, which is funded by NIH Office of Research Infrastructure Programs (P40 OD010440). This work was supported by the ShanghaiTech University start-up fund to Y.B.Q., the National Natural Science Foundation of China (32370601) to Y.B.Q., the National Natural Science Foundation of China (32370607) to Q.B., and the National Natural Science Foundation of China (32171156) to D.C.

## Methods

### Animals

Wild-type *Caenorhabditis elegans* were of the Bristol N2 strain. Worms were maintained on nematode growth medium (NGM) plates seeded with *E. coli* OP50 using standard methods. Unless otherwise noted, animals were cultured at 20 °C for experiments requiring precise developmental staging. All animals analyzed in this study were hermaphrodites.

### Allele and strain nomenclature

All alleles generated in the Yingchuan B. Qi laboratory are designated with the prefix ybq, and all strains are designated as BLW strains. Ex denotes extrachromosomal array transgenes, Is denotes integrated transgenes, and Si denotes single-copy integrated transgenes. The allele syb2564 was designed by Y.B.Q. and generated by SunyBiotech.

### Synchronization of worms

Gravid adult hermaphrodites were collected and bleached to isolate embryos. Eggs were washed at least three times with M9 buffer and transferred into 15 mL glass tubes. Embryos were allowed to hatch with gentle shaking in a 20 °C incubator to obtain synchronized L1 larvae. Synchronized animals were then transferred to NGM plates and cultured at 20 °C until the desired developmental stage.

### Microscopy, imaging, and quantification

Live animals were anesthetized with 0.5% (v/v) 1-phenoxy-2-propanol in M9 buffer and mounted on 3% agarose pads. Imaging was performed using an Olympus BX63 microscope. Confocal images were acquired on a Zeiss LSM 980 upright confocal microscope. Statistical analyses and graph generation were performed using GraphPad Prism.

### Fluorescence quantification

▪ **miRNA reporters and K10D3.4::mNG**: fluorescence intensity was measured in the head region and background-corrected using adjacent non-fluorescent areas.
▪ **DPY-4::mNG and DPY-5::mNG**: fluorescence intensity was measured across the worm body and background-corrected using adjacent regions.
▪ **NHR-23::mKate2, GFP::MYRF-1, and LIN-42::GFP**: nuclear fluorescence intensity was measured in head nuclei and background-corrected using adjacent areas. For nuclear penetrance analysis, animals with clearly detectable nuclear signals were scored as positive.
▪ **GFP::MYRF-1 in seam cells (NHR-23 AID experiments)**: images were processed by applying a Gaussian blur followed by subtraction of the blurred image from the original image to enhance signal-to-noise ratio.
▪ **GFP::MYRF-1 in *lin-42(0)* mutants**: fluorescence intensity was measured in the head region and background-corrected using adjacent areas.

### Molecular cloning

Repair templates for *lin-42* and *nhr-23* were amplified from the N2 genomic DNA. The lin-42::deg::GFP construct containing *loxP-unc-119(+)-loxP* in the third intron of GFP was commercially synthesized by Tsingke Biotechnology (Beijing, China), following the sequence described in Schwartz et al.^[52]^ The mKate2 fragment was amplified from plasmid pDD287, and mScarlet-I was amplified from VT3751 lysate. The AID2 cassette *[TIR1(F79G)-F2A-mTagBFP2-NLS-AID]*\* was modified from Ashley et al.^[38]^ Final constructs were assembled using the Vazyme ClonExpress Ultra One-Step Cloning Kit V2 (C116).

### CRISPR–Cas9 genome editing

CRISPR–Cas9 genome editing was performed using plasmid-based Cas9 and sgRNA expression. Two sgRNAs were used for each knock-in strain. Individual F1 progeny from microinjected animals were screened by PCR.

**nhr-23::mKate2(ybq252)**

sgRNA1: ttgctccagaataagtccagcgg

sgRNA2: cttattgtacaaataaccggagg

nhr-23 WT(background):

…gtctgatccaacatcatctgaaaagcttcctgccctctacaaagagctattcactgcagatcggcct-[insertion point]-tgactgaatccatatatcatcaatagttttatccatgctctccttccctatccccgt…

nhr-23::mKate2(yba252):

…gtctgatccaacatcatctgaaaagcttcctgccctctacaaagagctattcactgcagatcggcct-[mKate2]-tgactgaatccatatatcatcaatagttttatccatgctctccttccctatccccgt…

**lin-42::m-Scarlet-I(ybq251)**

sgRNA1: gaagttttcgggaagacgggcgg

sgRNA2: tgtttttctacatgactcctcggcgg

lin-42 WT(background):

…aaaacttcctcatcctcctctctcctaatgctacgggattctcagaat-[insertion point]-taagctactgcccaatttttctttcaatttttaatttttttcgaa…

lin-42::m-Scarlet-I(ybq251):

…aaaacttcctcatcctcctctctcctaatgctacgggattctcagaat-[mScarlat-I]-taagctactgcccaatttttctttcaatttttaatttttttcgaa…

**lin-42::deg::GFP(ybq224)**

This allele was generated in strain EG9881 as described previously^[52]^. UNC-119–rescued F1 animals were isolated, and the *loxP-unc-119(+)-loxP* cassette was removed by heat-shock–induced Cre recombination.

sgRNA1: gaagttttcgggaagacgggcgg

sgRNA2: tgtttttctacatgactcctcggcgg

lin-42 WT(background):

…aaaacttcctcatcctcctctctcctaatgctacgggattctcagaat-[insertion point]-taagctactgcccaatttttctttcaatttttaatttttttcgaa…

lin-42::deg::GFP(ybq224):

…aaaacttcctcatcctcctctctcctaatgctacgggattctcagaat-[deg-GFP]-taagctactgcccaatttttctttcaatttttaatttttttcgaa…

**GFP::myrf-1 (syb2564)**

This allele was generated by SunyBiotech. GFP was inserted at the N-terminus of MYRF-1.

sgRNA1: cacttcgaacATGTCGAGCTCGG

sgRNA2: TCGAGCTCGGACCTTCTGAAAGg

myrf-1 WT (background):

…aagccctcccacatacccaataccaaagaaccgatactttagcacacttcgaacATG-[insertion point]-TCGAGCTCGGACCTT<u>CTG</u>AAAGgtaagacatattaaaaaaaaataggttttgacggt

GFP::myrf-1(syb2564):

…aagccctcccacatacccaataccaaagaaccgatactttagcacacttcgaacATG-[GFP]-

-[GGTGGCGGA]-

TCGAGCTCGGACCTTCTCAAAGgtaagacatattaaaaaaaaataggttttgacggt

### Single-copy and extrachromosomal transgene alleles

Single-copy transgene alleles were generated using the miniMOS system as described previously^[53]^. Injection mixtures containing Mos1 transposase, fluorescent coinjection markers, and the corresponding miniMOS plasmids were microinjected into the gonads of young adult hermaphrodites. Transgenic lines were selected using antibiotic resistance (hygromycin or G418, as appropriate), isolated after 7–10 days, and outcrossed to N2.

The AID2 allele ybqSi295 [Peft−3::TIR1(F79G)−F2A−mTagBFP2−NLS−AID∗] was generated by injecting pQA2426 together with pCFJ601 (Mos1 transposase) and pCFJ104 (Pmyo-3-mCherry) into BLW2652, followed by hygromycin selection.

The epidermal and seam cell membrane markers ybqSi306 [dpy−5p::TagRFP::PH+NeoR] and ybqSi296 [Pscm::TagRFP::PH+NeoR] were generated using similar miniMOS injection strategies with G418 selection in BLW1827 and BLW889 backgrounds, respectively.

The extrachromosomal array ybqEx962 [Pvha−6::lin−42a] was generated by injecting pQA2428 together with pCFJ104 (Pmyo-3-mCherry) into BLW889, and F1 animals carrying arrays were identified by fluorescence.

### Auxin-inducible degradation (AID)

Stock solutions were prepared as follows: **AID1**: 50 mM K-NAA (potassium 1-naphthaleneacetate) in ddH₂O; **AID2**: 5 mM 5-Ph-IAA in DMSO. Final concentrations were 4 mM K-NAA for AID1 plates and 5 µM 5-Ph-IAA for AID2 plates. For AID, stock solutions were first diluted in ddH₂O before addition to NGM with OP50. Plates were air-dried in the dark and stored until use.

### RNA interference

dsRNA expression was induced with 0.4 mM IPTG. For seam-cell MYRF-1 RNAi screening, NGM plates containing 0.4 mM IPTG and 4 mM K-NAA were used. Three L4 hermaphrodites were placed on each plate and cultured for 60 h before phenotype scoring.

### RNA-seq preparation and analysis

Synchronized animals were cultured on control or AID2 plates and harvested at 15.5 h and 25.5 h post-hatching. For each condition and time point, >2,000 animals were collected per sample, with three biological replicates. Worms were washed with M9, pelleted, resuspended in 100 µL M9, mixed with an equal volume of TRIzol, flash-frozen in liquid nitrogen, cryogenically ground, and homogenized in 1 mL TRIzol for RNA extraction.

Library preparation, sequencing, and primary analysis were performed by Novogene (Beijing, China). TPM values were calculated based on gene length and mapped read counts. Differential expression analysis was conducted using DESeq2 (v1.42.0), with Benjamini–Hochberg correction (padj ≤ 0.05, |log₂FC| ≥ 1).

GO enrichment was performed using ClusterProfiler (v4.8.1), with gene-length bias correction. Enrichment P values were calculated using a one-tailed hypergeometric test followed by Benjamini–Hochberg multiple-testing correction. The top 10 enriched GO terms in the Biological Process (BP), Molecular Function (MF), and Cellular Component (CC) categories, ranked by P values, were visualized using dot plots. Venn diagrams were generated using eulerr, volcano plots using EnhancedVolcano, and oscillatory gene heatmaps using pheatmap.

### MYRF-1 (PQN-47) ChIP–seq

MYRF-1 ChIP–seq was performed through the modENCODE/modERN^[47,54]^pipeline using a GFP-tagged MYRF-1 strain (*GFP::myrf-1*, syb2564). Worms were synchronized at L1 and collected at ∼27 h post-hatching (late L2). Chromatin was immunoprecipitated using an in-house goat anti-GFP antibody, with matched input controls. Library preparation, sequencing, mapping, peak calling, and QC followed standardized modERN workflows. Processed datasets are available through the modERN resource portal: modERN resource portal (https://epic.gs.washington.edu/modERNresource/).

### ChIP–seq annotation and GO analysis

Optimal MYRF-1 ChIP–seq peaks were identified through the standardized modERN ChIP–seq pipeline. Genomic annotation and TSS-centric visualization were performed using ChIPseeker (v1.30.2). GO enrichment of MYRF-1–bound genes was performed using ClusterProfiler (v4.7.1.003), with Benjamini–Hochberg correction. The top 10 GO terms in BP, MF, and CC categories were visualized using dot plots.

## Author Contributions

Z.W. and S.S. performed the majority of experiments and data analysis. X.F. contributed to the analysis of *dpy-5p* expression and molting phenotypes. D.C. assisted with the development and optimization of AID reagents. Q.B. performed bioinformatic analyses of ChIP–seq and RNA-seq datasets. Z.W. conceived the AID experiments and contributed to manuscript drafting. Y.B.Q. conceived and supervised the project, directed the research, and wrote the manuscript. D.C., Q.B., and Y.B.Q. acquired funding for the study.

---

**Figure S1.**
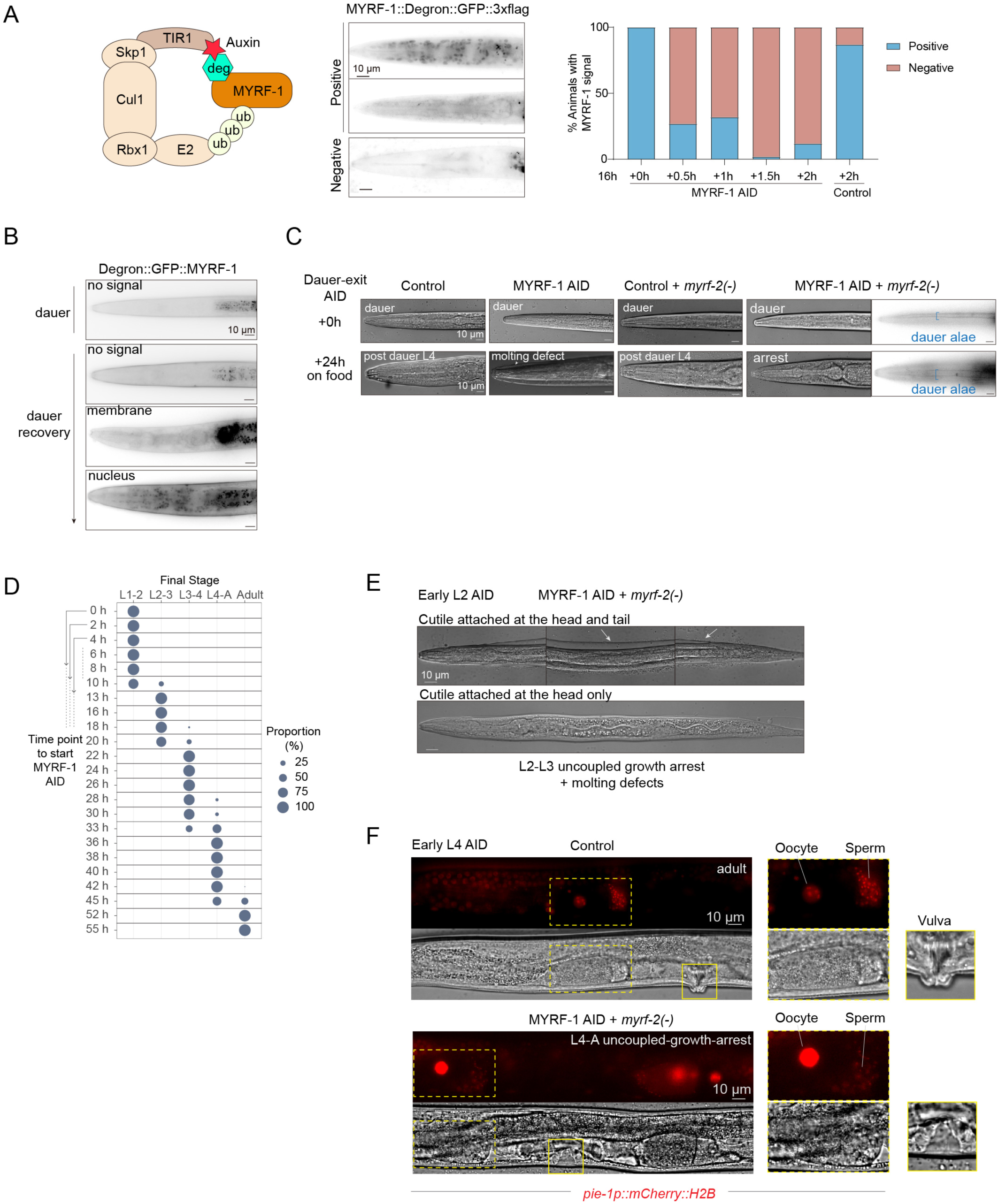
Validation and developmental consequences of MYRF-1 auxin-induced degradation (AID). **A.** Schematic of the auxin-inducible degron system used to acutely deplete MYRF-1. Representative images of MYRF-1::Degron::GFP::3xFLAG showing efficient loss of MYRF-1 signal upon auxin addition, with quantification of the fraction of animals positive or negative for MYRF-1 signal over time. **B.** Localization of Degron::GFP::MYRF-1 in dauer and during dauer recovery. MYRF-1 signal is absent in dauer larvae but reappears at the membrane and subsequently in nuclei during recovery. **C.** Developmental outcomes following dauer recovery with or without MYRF-1 AID. MYRF-1 depletion alone results in molting defects or arrest after dauer exit. MYRF-1 AID; *myrf-2(–)* causes uncoupled growth arrest prior to the dauer–L4 molt, accompanied by persistence of dauer alae. **D.** Stage-specific outcomes following MYRF-1 AID initiated at time point series after hatching. Bubble plot indicates the proportion of animals arresting at specific stages depending on the timing of MYRF-1 depletion. **E.** Representative differential interference contrast (DIC) images showing incomplete L2–L3 molting in MYRF-1 AID; *myrf-2(–)* animals, with unshed cuticle remaining attached at the head and/or tail. **F.** Early L4–initiated MYRF-1 AID in a *myrf-2(–)* background results in arrest of vulval morphogenesis at mid-L4, whereas control animals form an adult vulva. In contrast, gonadal development is largely unaffected by *myrf-1/2* deficiency, with continued germline progression and the presence of oocytes and sperm, visualized using a germline-expressed nuclear red histone marker.

**Figure S2.**
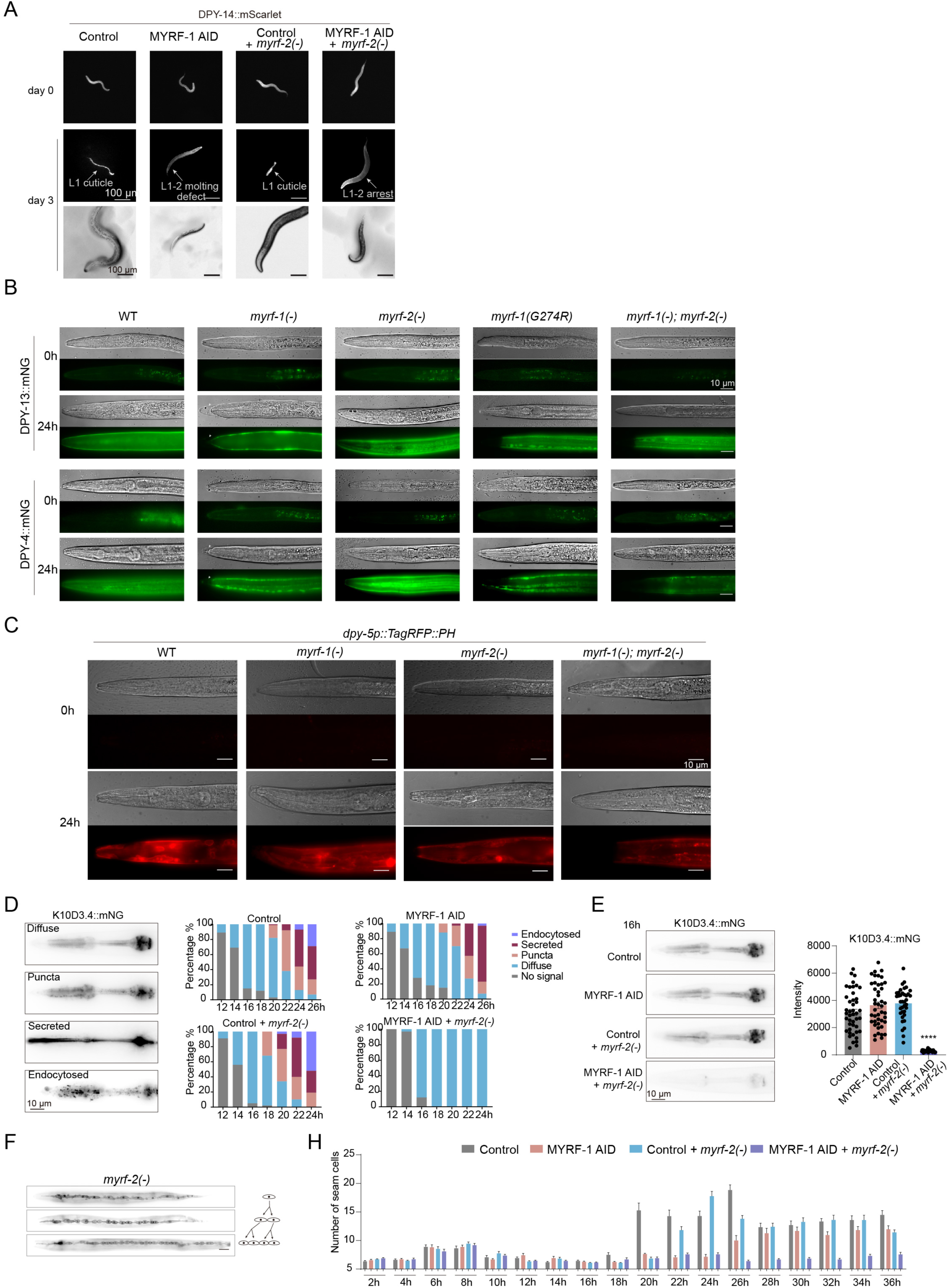
Additional analyses of cuticle remodeling, molting, and seam cell behavior in MYRF-deficient animals. **A.** Tracking individual animals using DPY-14::mScarlet to label the L1 cuticle and monitor L1 cuticle shedding. Control and *myrf-2(–)* animals shed the DPY-14–marked L1 cuticle on plates, whereas MYRF-1 AID animals display L1–L2 molting defects with retention of unshed DPY-14–positive cuticle. MYRF-1 AID; *myrf-2(–)* animals arrest while carrying persistent DPY-14–positive cuticles. **B.** Expression of the L2 cuticle components DPY-13::mNG and DPY-4::mNG at 0 h and 24 h. WT, *myrf-1(–)*, and *myrf-2(–)* animals express both markers. *myrf-1(G274R)* and *myrf-1(–); myrf-2(–)* animals also express DPY-13::mNG and DPY-4::mNG, but reporter signals are absent from the head region. Notably, in *myrf-1(–)* and *myrf-1(G274R)* mutants, the reporters label the newly synthesized, internal cuticle rather than the unshed external cuticle. **C.** Expression of the transcriptional reporter *dpy-5p::TagRFP::PH*. The reporter is expressed in *myrf-1(–); myrf-2(–)* animals but is absent from the head region. **D.** Subcellular localization patterns of the secreted cuticle-associated protein K10D3.4::mNG (diffuse, punctate, secreted, endocytosed) over time in control, MYRF-1 AID, *myrf-2(–)*, and MYRF-1 AID; *myrf-2(–)* animals, with quantification of localization categories. **E.** Representative images and fluorescence quantification of K10D3.4::mNG showing strong reduction of signal in MYRF-1 AID; *myrf-2(–)* animals. **F.** Representative images of seam cell morphology in *myrf-2(–)* animals. **G.** Quantification of seam cell number over time. MYRF-1 AID animals undergo two rounds of seam cell division, whereas MYRF-1 AID; *myrf-2(–)* animals fail to undergo seam cell divisions.

**Figure S3.**
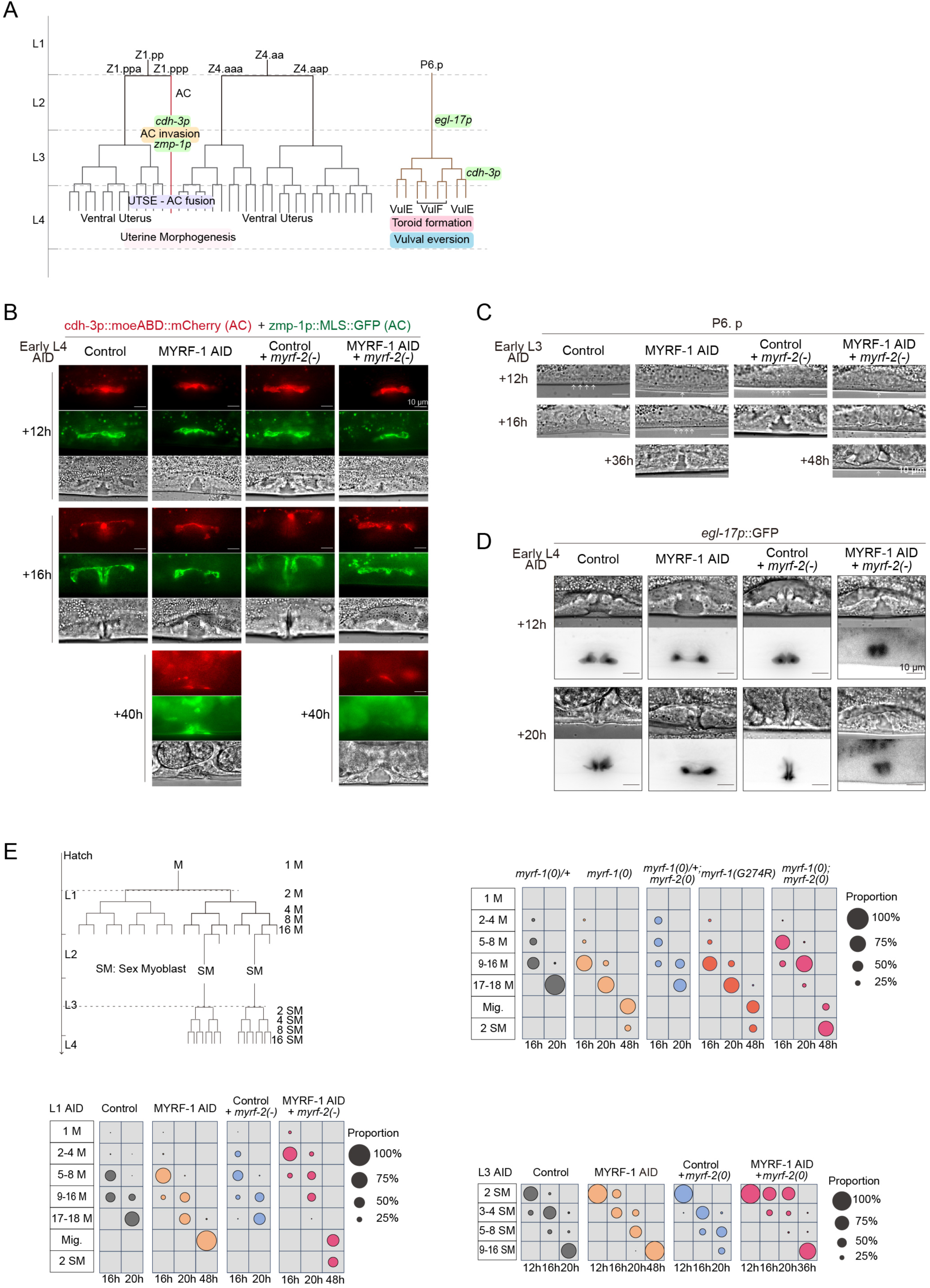
MYRF is required for proper progression of uterine and vulval development, but is largely dispensable for the M-lineage program. **A.** Schematic of anchor cell (AC), uterine, and vulval lineages, highlighting key morphogenetic events and marker expression used in this study, including AC invasion, AC–utse fusion, and vulval toroid formation and eversion. **B.** AC–UTSE fusion visualized using cdh-3p and zmp-1p markers. In early L4, AC–UTSE fusion occurs in both MYRF-1 AID and MYRF-1 AID; *myrf-2*(–) animals, consistent with the fact that this event normally precedes late L4. In contrast, vulval morphogenesis is delayed and morphologically abnormal in MYRF-1 AID animals and is completely arrested at a mid-L4–like state in MYRF-1 AID; *myrf-2*(–) animals. Although cdh-3p and zmp-1p are normally activated later in vulval lineages, this vulval expression is absent in MYRF-1 AID; *myrf-2*(–) animals. **C.** Vulval precursor cell (P6.p) divisions following early L3 MYRF-1 AID. Representative DIC images show delayed vulval cell divisions in MYRF-1 AID animals, whereas MYRF-1 AID; *myrf-2*(–) animals exhibit a non-dividing P6.p lineage. **D.** Expression of the vulval fate marker egl-17p::GFP after early L4 MYRF-1 AID. MYRF-1 AID animals initiate vulval development but display defective morphogenesis, whereas MYRF-1 AID; *myrf-2*(–) animals completely fail to initiate vulval morphogenesis. **E.** Lineage schematic of post-embryonic M-cell divisions (left) and quantitative analysis of M-lineage progression under the indicated genotypes and MYRF-1 AID conditions (right and below). Bubble size indicates the proportion of animals reaching each division stage at the specified time points. M-cell divisions proceed largely normally in *myrf* mutants and under MYRF-1 AID conditions. Sex myoblast (SM) divisions are delayed but largely completed in MYRF-1 AID animals.

**Figure S4.**
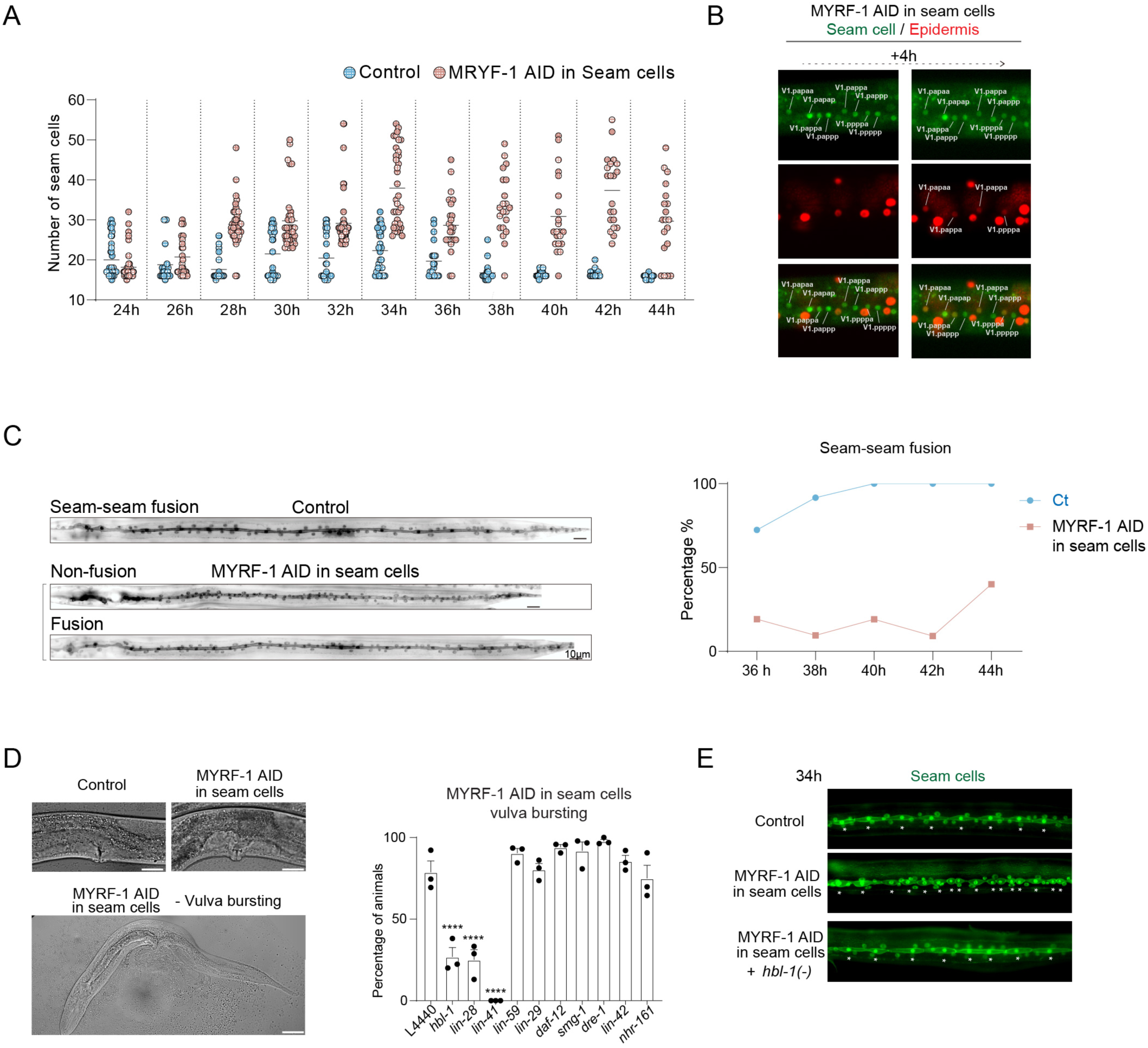
Seam cell–specific MYRF-1 depletion causes heterochronic division defects and vulval bursting. **A.** Quantification of seam cell number over time in control animals and animals with seam cell–specific MYRF-1 AID. MYRF-1 depletion in seam cells leads to excess seam cell accumulation beginning in L3, consistent with reiteration of L2-like divisions. **B.** Live imaging of the same animals (L3) tracking epidermal fusion of differentiated seam cell daughter cells generated by reiterated L2-like divisions. Images are shown at 4-h intervals. **C.** Analysis of final seam-seam fusion during late larval stages. Left, representative images showing fused and unfused differentiated seam cell daughters. Right, quantification of the percentage of animals exhibiting seam-seam fusion over time. Seam cell–specific MYRF-1 AID delays or prevents epidermal fusion. **D.** Seam cell–specific MYRF-1 AID causes highly penetrant vulval bursting. Left, representative DIC images. Right, quantification of vulval bursting in control and after RNAi knockdown of heterochronic pathway components. RNAi of *hbl-1*, *lin-28*, and *lin-41* significantly suppresses bursting. **E.** Representative animals showing that the increased seam cell number caused by seam cell–specific MYRF-1 AID is suppressed by *hbl-1* loss of function.

**Figure S5.**
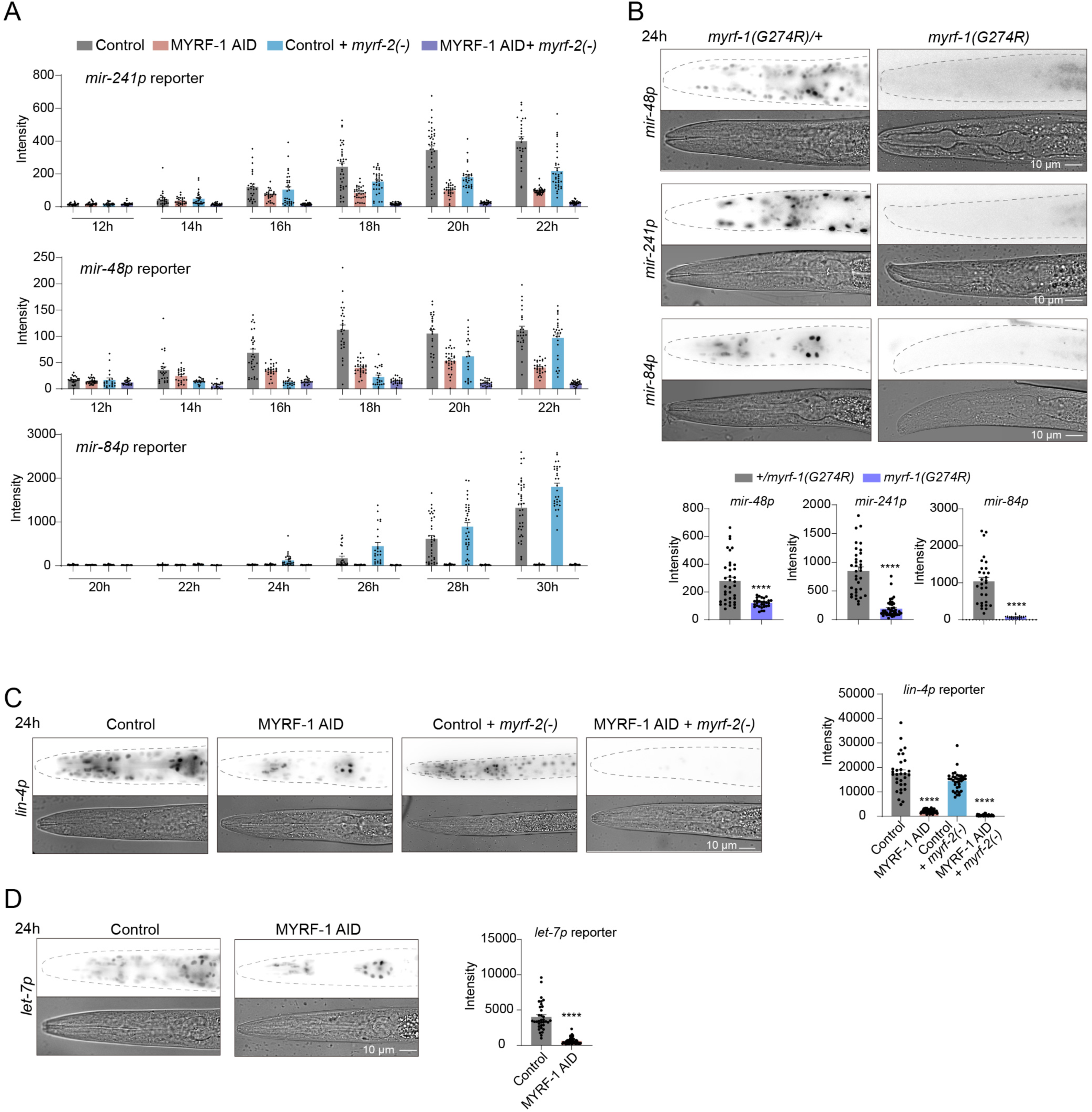
MYRF activity is required for expression of heterochronic microRNAs. **A.** Quantification of *mir-241p*, *mir-48p*, and *mir-84p* reporter intensities over time in control and MYRF-1 AID conditions, revealing the severe suppression of the reporter expression by MYRF-1 AID; *myrf-2*(–). **B.** Representative images of *mir-48p*, *mir-241p*, and *mir-84p* reporters in heterozygous and homozygous *myrf-1(G274R)* animals, indicating reduced reporter expression in the mutant background. Below: Quantification of reporter intensities. **C.** Expression of the *lin-4p* reporter in control and MYRF-1 AID conditions, showing strong dependence on MYRF activity. **D.** *let-7p* reporter expression in control and MYRF-1 AID animals, demonstrating reduced transcriptional activation upon MYRF depletion.

**Figure S6.**
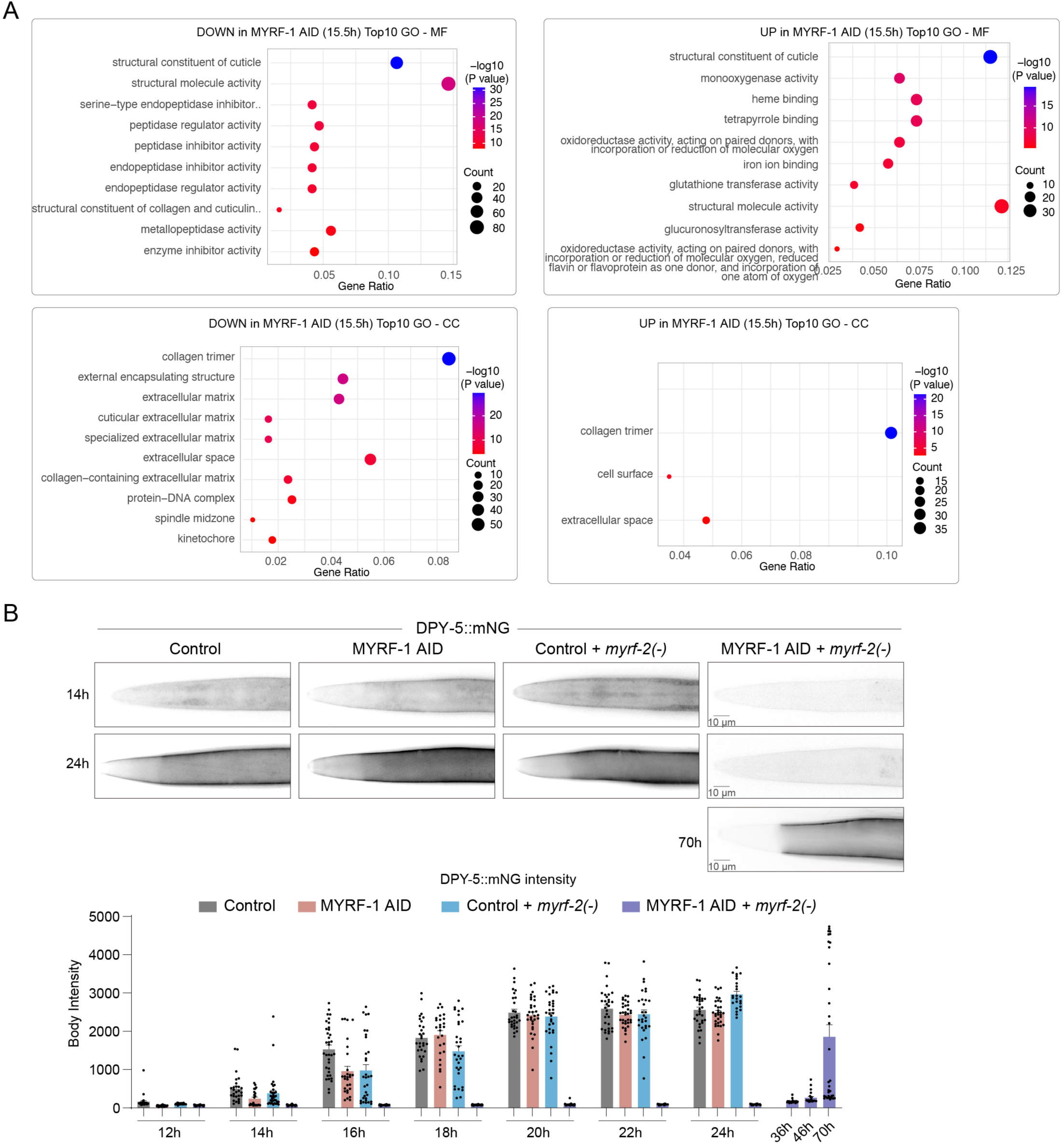
MYRF depletion alters molting-associated gene expression and cuticle protein dynamics. **A.** Gene Ontology (GO) enrichment analysis of molecular function (MF) and cellular component (CC) categories for genes downregulated (left) or upregulated (right) following MYRF-1 AID at 15.5 h. Downregulated genes are enriched for cuticle structural components, extracellular matrix, and protease inhibitor activities, whereas upregulated genes are enriched for metabolic and detoxification-related functions. **B.** Analysis of the L2 cuticle collagen reporter DPY-5::mNG. Representative images (top) and quantitative fluorescence measurements (bottom) show expression of DPY-5 in control and *myrf-2*(–) animals, reduced or delayed expression upon MYRF-1 AID, and minimal activation in MYRF-1 AID + *myrf-2*(–) animals, despite gradual protein accumulation in arrested animals.

**Figure S7.**
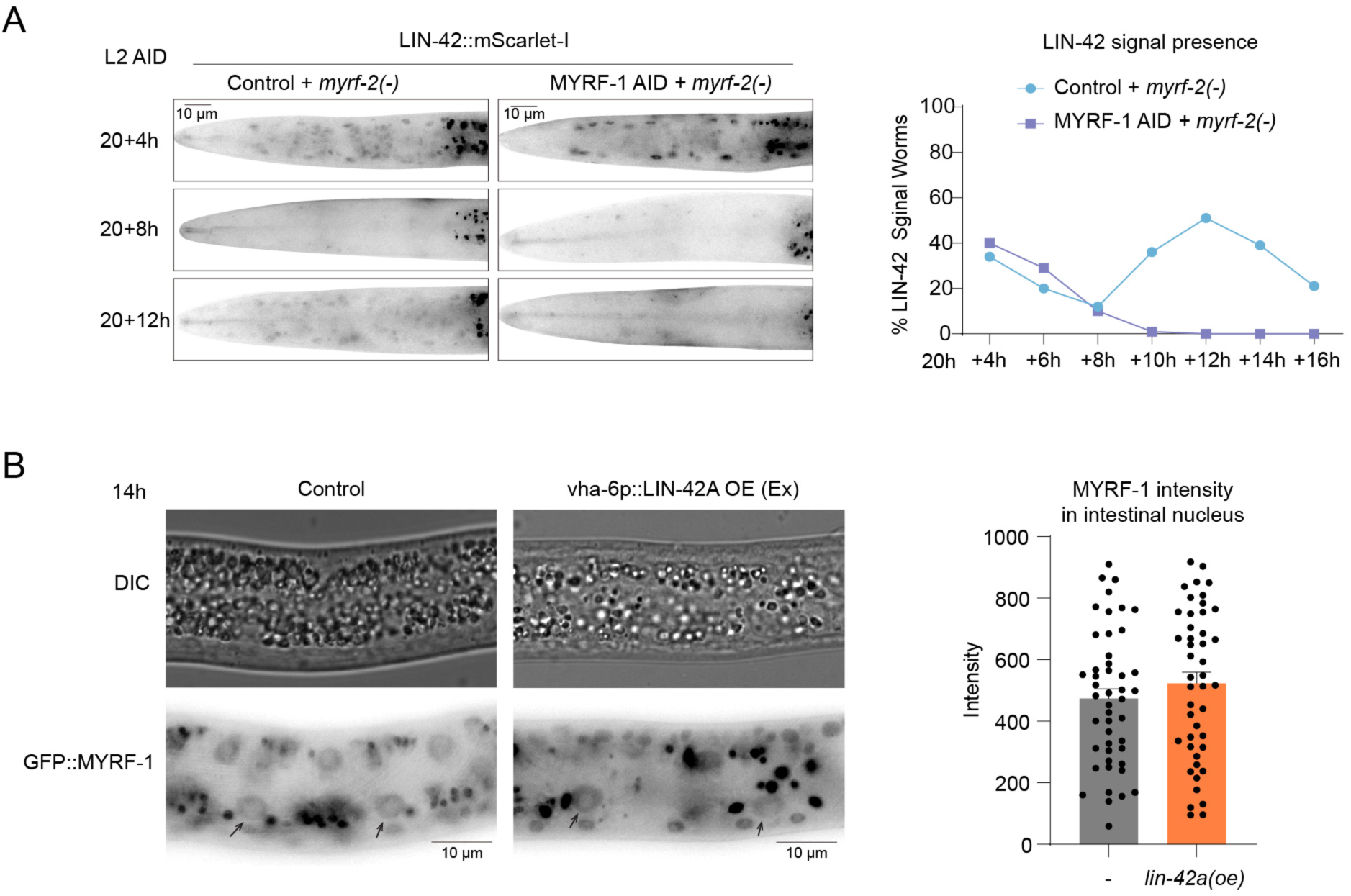
Analysis of functional interactions between MYRF-1 and LIN-42. **A.** Analysis of LIN-42::mScarlet-I signal following L2-stage MYRF-1 AID in a *myrf-2*(–) background. Left, representative images showing reappearance of LIN-42 expression at late L2 in control animals but suppression of LIN-42 signal in MYRF-1 AID conditions. Right, quantification of the fraction of animals with detectable LIN-42 signal over time. **B.** Effect of LIN-42 overexpression on MYRF-1 nuclear accumulation. Representative DIC and GFP images show comparable GFP::MYRF-1 nuclear localization in intestinal cells of control animals and animals overexpressing LIN-42A (*vha-6p::lin-42a*). Quantification (right) shows no significant difference in nuclear GFP::MYRF-1.

## Supplementary Files

- **File S1. MYRF-1 ChIP-seq peaks mapped to microRNA loci** Excel file listing MYRF-1 ChIP-seq peaks mapped to microRNA genes, including peak coordinates, genomic annotation, and sorted by peak ranks. Note that peaks in this list are assigned based on proximity to annotated microRNA genes; some peaks may also lie near or overlap protein-coding genes, and this file does not distinguish between these assignments.
- **File S2. MYRF-1 ChIP-seq peaks mapped to protein-coding genes** Excel file listing MYRF-1 ChIP-seq peaks associated with protein-coding genes, including peak coordinates, genomic annotations, and ranking by peak strength. Peaks are annotated as intragenic or intergenic–upstream relative to the nearest gene; intergenic–downstream peaks are excluded from this list.
- **File S3. Gene Ontology enrichment of MYRF-1 ChIP-seq target genes** Excel file summarizing GO enrichment analysis of protein-coding genes bound by MYRF-1, including Biological Process (BP), Molecular Function (MF), and Cellular Component (CC) categories, enrichment statistics, adjusted P values, and gene IDs.
- **File S4. Differentially expressed genes following MYRF-1 AID at 15.5 h** Excel file containing RNA-seq differential expression results comparing control and MYRF-1 AID; myrf-2(-) animals at 15.5 h post-hatching, including log2 fold change, adjusted P values, and FPKM values. Differentially expressed genes (DEGs) are organized into three separate sheets, comprising the full DEG list, upregulated genes, and downregulated genes.
- **File S5. Gene Ontology enrichment of MYRF-1 AID-regulated genes at 15.5 h** Excel file reporting Gene Ontology (GO) enrichment analysis of genes upregulated or downregulated following MYRF-1 depletion at 15.5 h, including enrichment statistics for Biological Process (BP), Molecular Function (MF), and Cellular Component (CC) categories. Results are organized into six separate sheets: Down-BP, Down-MF, Down-CC, Up-BP, Up-MF, and Up-CC.
- **File S6. Oscillatory genes affected by MYRF-1 AID at 15.5 h** Excel file identifying oscillatory genes (based on published larval oscillation datasets) that are differentially expressed upon MYRF depletion at 15.5 h, including expression changes.
- **File S7. C. elegans strains used in this study** Excel file listing all strains used, including strain number, genotype and allele information.

